# Strand swapping in the LolA-like protein GerS promotes the allosteric activation of a bacterial amidase in *Clostridioides difficile*

**DOI:** 10.1101/2025.01.13.632879

**Authors:** Jacob M. Bouchier, Aimee Shen

## Abstract

GerS is a key lipoprotein regulator of *Clostridioides difficile* spore germination that shares structural, but not sequence, similarity to LolA in Gram-negative bacteria. GerS and LolA have vastly different biological roles: LolA functions in the trafficking of lipoproteins across the periplasmic space to the outer membrane of Gram-negative bacteria, while GerS allosterically activates the germination-specific amidase CwlD in endospore-forming Gram-positive bacteria. Here, we explore the diversity of LolA-like proteins across bacteria and reveal that GerS homodimer formation is a conserved feature of Peptostreptococcaceae family orthologs. Since dimerization of *C. difficile* GerS occurs via strand-swapping, we sought to investigate the functional importance of its strand swap by identifying mutations that alter the GerS dimer:monomer equilibrium. Our analyses reveal that GerS^K71R^ and GerS^N74L^ substitutions in the strand-swap hinge of GerS stabilize the homodimer, while a GerS^T72P^ substitution promotes monomer formation. Conversely, substituting the highly conserved proline at the equivalent site in LolA for threonine converts LolA from a monomer to a homodimer. Since our data indicate that destabilizing the GerS dimer impairs both GerS:CwlD binding *in vitro* and CwlD function in *C. difficile*, strand-swapped dimer formation in LolA-like proteins enhances their affinity for their binding partners. These data are consistent with analyses of equivalent LolA mutants in *E. coli* and thus provide new insight into the evolution and specialization of this intriguing group of proteins across the bacterial domain.

## Introduction

The InterPro homologous superfamily “Prokaryotic lipoproteins and lipoprotein localization factors” (PLLLF; SSF89392) encompasses a group of structurally similar proteins that are widely distributed across the bacterial domain. While some proteins of this group function as anti-sigma factors [1,2], most function as chaperones for transporting proteins or glycolipids across the periplasmic space of diderm bacteria. For example, LolA, the archetypal PLLLF protein, functions within the Lol system (LolABCDE) to facilitate lipoprotein trafficking across the periplasm in diderm bacteria [3–9], while LppX [10], LprG [11], and LprF [12] transport various glycolipids across the periplasm of *Mycobacterium* spp. In contrast, PLLLF proteins in endospore-forming monoderm bacteria seem to have evolved specialized roles in sporulation and germination. For example, SsdC from *Bacillus subtilis* forms a ring-like structure in the forespore that functions in the development of spore shape [13], and GerS regulates spore germination in *Clostridioides difficile* [14].

Spore germination depends on the enzymatic degradation of the cortex, a protective layer of modified peptidoglycan that maintains metabolic dormancy [15,16]. The cortex lytic enzymes that degrade the cortex recognize spore-specific peptidoglycan modifications found exclusively within the cortex, namely muramyl-δ-lactam (MAL) [17]. The specificity of these enzymes for MAL limits their degradative activities to the cortex layer. MAL is made by the coordinated activities of the germination-specific amidase CwlD and the peptidoglycan deacetylase PdaA [18–21]. While *B. subtilis* CwlD has intrinsic activity, *C. difficile* CwlD requires an additional factor to license its activity, the GerS lipoprotein [22,23]. GerS forms a 1:1 complex with CwlD and allosterically activates CwlD’s amidase activity by stabilizing CwlD’s binding to its Zn^2+^ cofactor [23]. This interaction is essential for germination because deletion of *gerS* abolishes spore germination in *C. difficile* [14] and disruption of a salt bridge formed between aspartate 106 of GerS and arginine 169 of CwlD reduces germination [23]. However, the reduction in *C. difficile* spore germination caused by the GerS D106R substitution is relatively subtle, even though it significantly reduces binding to CwlD *in vitro* [23]. These data indicate that additional factors beyond the salt bridge regulate the GerS:CwlD interaction and thus the amidase activity of CwlD.

In the crystal structure of the GerS:CwlD complex, GerS forms a homodimer via a strand-swap mechanism [23]. Strand swapping, a specific form of domain swapping [24–31], is a dimerization mechanism in which one or more β-strands are exchanged between two proteins [32]. Dimerization via strand swapping is observed in the dimeric structures found in the light chain subunits of immunoglobulins [33] and cystatins [34], and it is thought to be associated with pathogenic amyloid fibril formation in humans [35–37]. Strand swapping can also regulate protein function [38]. For example, monomeric human cystatin C inhibits cathepsin B, but its strand-swapped dimer loses this ability [39], and strand swapping in cadherins facilitates cell-cell adhesion, forming weak, reversible interactions [28,40]. However, the functional significance of strand swapping in GerS on the allosteric activation of CwlD by GerS remains unclear.

Notably, *C. difficile* GerS is the only known PLLLF protein that naturally dimerizes via a strand swap mechanism, whereas LolA from *Escherichia coli* can be induced to dimerize when Phe68 is substituted with Glu, which induces a similar strand swap [41,42]; the F68E substitution was designated F47E based on the sequence after lipoprotein signal peptide processing. This mutation is toxic to *E. coli* because LolA^F68E^ binds LolC too tightly and prevents proper lipoprotein trafficking to the outer membrane [42]. Here, we test the impact of strand swapping in GerS on CwlD function. By identifying mutations that shift the GerS dimer:monomer equilibrium, we demonstrate that strand-swapping in GerS stabilizes its interaction with CwlD and promotes the allosteric activation of this important amidase. Our analyses provide molecular insight into the mechanisms that allow for strand swapping and highlight the general role of these structures in regulating adhesive interactions.

## Results

### PLLLF proteins maintain low sequence identity but high sequence similarity and structural similarity

With the goal of identifying the underlying and shared features between GerS and the rest of the PLLLF superfamily, we analyzed the evolutionary relatedness of proteins within the group. We found that heuristic search methods, such as BLASTp, were inadequate for identifying members of this group. Instead, we employed probabilistic searches using InterPro and structural searches using FoldSeek to identify which major bacterial classes contain bacteria that encode PLLLF proteins [43–46]. We then generated a phylogenetic tree using core genes across species based on the major bacterial classes that have species encoding PLLLF proteins (Supplemental Figure 1). Remarkably, we found that PLLLF proteins are widespread across the bacterial domain. Further, we found that the presence of PLLLF proteins is independent of whether the bacteria are monoderms or diderms (Supplemental Figure 1), suggesting that not all PLLLF proteins function in a similar way to LolA.

Using AlphaFold 3 [47], we predicted the structure of 17 representative PLLLF proteins and aligned them using DALI [48]. Using this structural alignment, we compared sequence identity and sequence similarity across the proteins. Despite sharing little sequence identity, the sequence similarity across PLLLF group members is high (Figure 1A). Together, these observations indicate that PLLLF proteins are highly conserved and suggest that they share evolutionary ancestry, although they likely evolved specialized roles within diderms vs. monoderms.

**Figure 1.**
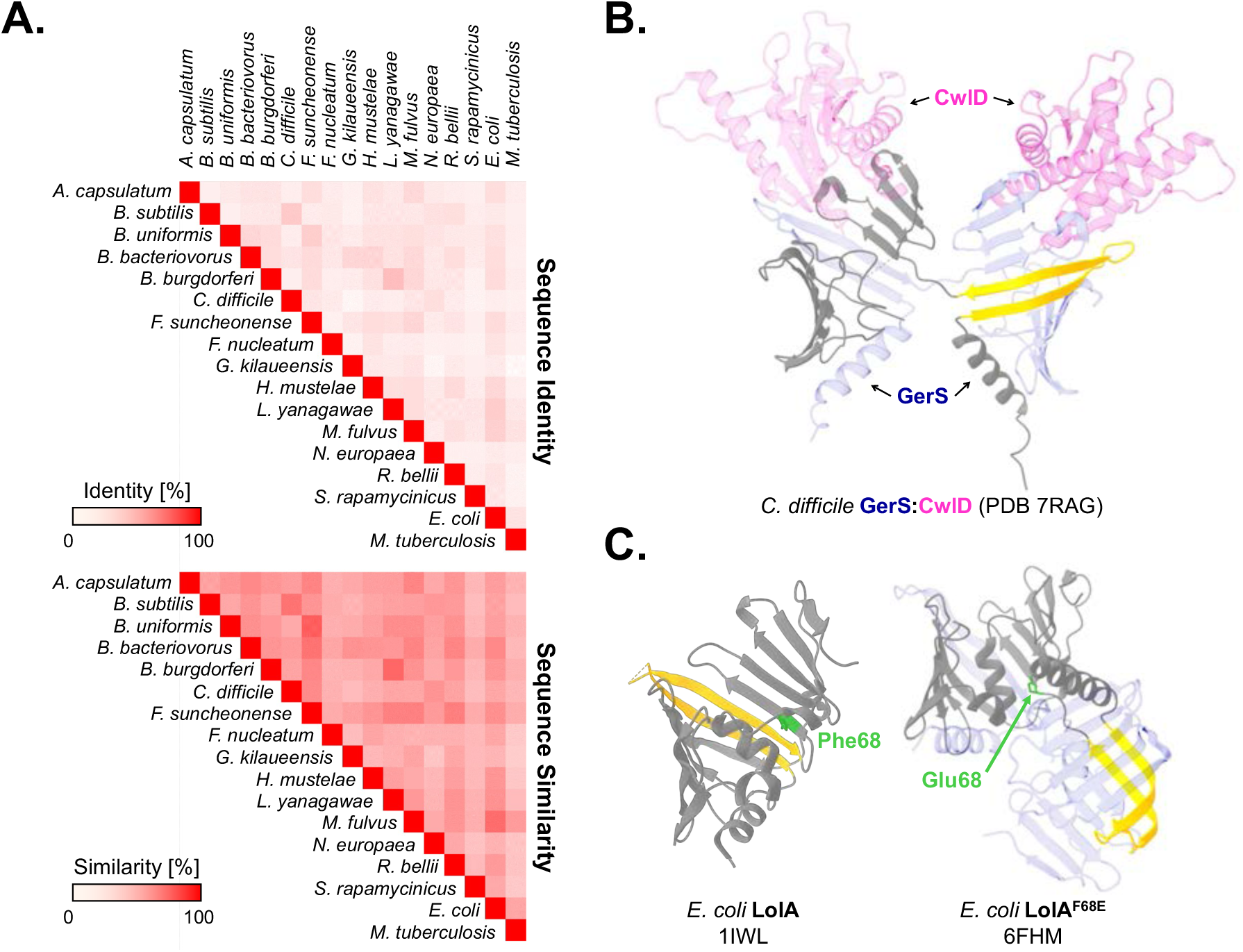
PLLLF proteins are structurally similar but share low sequence identity. A) Comparisons of sequence identity and sequence similarity between 17 representative PLLLF proteins. Gradients represent percentages from 0 to 100. B) Ribbon representation of the crystal structure of the GerS-CwlD complex (PBD 7RAG) with the GerS strand swap colored yellow. The other GerS is colored in transparent blue, and both CwlD proteins are colored in transparent pink. C) Ribbon representation of the crystal structures of the *Escherichia coli* LolA^WT^ (PDB 1IWL) and LolA^F68E^ (PDB 6FHM). The strand swap in LolA^F68E^ structure is shown in yellow; the corresponding β-strand in LolA^WT^ is shown in yellow. The other LolA^F68E^ protomer in the homodimer is colored in transparent blue. Residue 68, which is a phenylalanine in LolA^WT^ and a glutamate in LolA^F68E^, is colored green.

### GerS strand swapping is unique among characterized PLLLF proteins

While AlphaFold 3 [47] predicts that GerS adopts a structure remarkably similar to other PLLLF proteins, the GerS-CwlD crystal structure shows that GerS forms a homodimer via a strand-swap mechanism (Figure 1B)[23]. Crystal structures of other PLLLF proteins, ranging across several bacterial classes, suggest that PLLLF proteins exist either as monomers or heterodimers with other proteins but rarely as homodimers [3,5,10,11,49,50].

While LolA does not form homodimers naturally, a single substitution of phenylalanine to glutamate at residue 68 (F68E, previously published as F47E based on numbering from the lipobox) is sufficient to induce homodimerization [42]. This homodimerization is achieved by converting a β-turn into a strand-swap hinge, which is remarkably similar to the strand swap observed in GerS (Figure 1C, Supplemental Figure 2). Notably, the LolA^F68E^ variant binds its interacting partner LolC with increased affinity, but this enhanced affinity interferes with LolA’s ability to bind the lipoprotein substrates that it normally transports across the periplasm, rendering the LolA^F68E^ variant toxic to *E. coli* [41,42]. Based on these observations, we hypothesized that strand swap-mediated dimerization of GerS stabilizes GerS binding to CwlD and promotes CwlD binding to its Zn^2+^ co-factor.

### Dimerization is conserved in GerS orthologs of the Peptostreptococcaceae family

Since CwlD is widely conserved in endospore-forming bacteria, we first assessed the conservation of *gerS* among the Firmicutes, the only known class to contain endospore-forming bacteria like *C. difficile* and *Bacillus subtilis*. To avoid issues associated with identifying PLLLF proteins through heuristic methods like BLASTp, we instead used synteny-based searches such as GeCoViz [51] and BioCyc genome alignment [52] to find GerS orthologs; we confirmed their structural homology using AlphaFold3-predicted structures and ChimeraX (Figure 2A; Supplemental Figure 3) [47,53].

**Figure 2.**
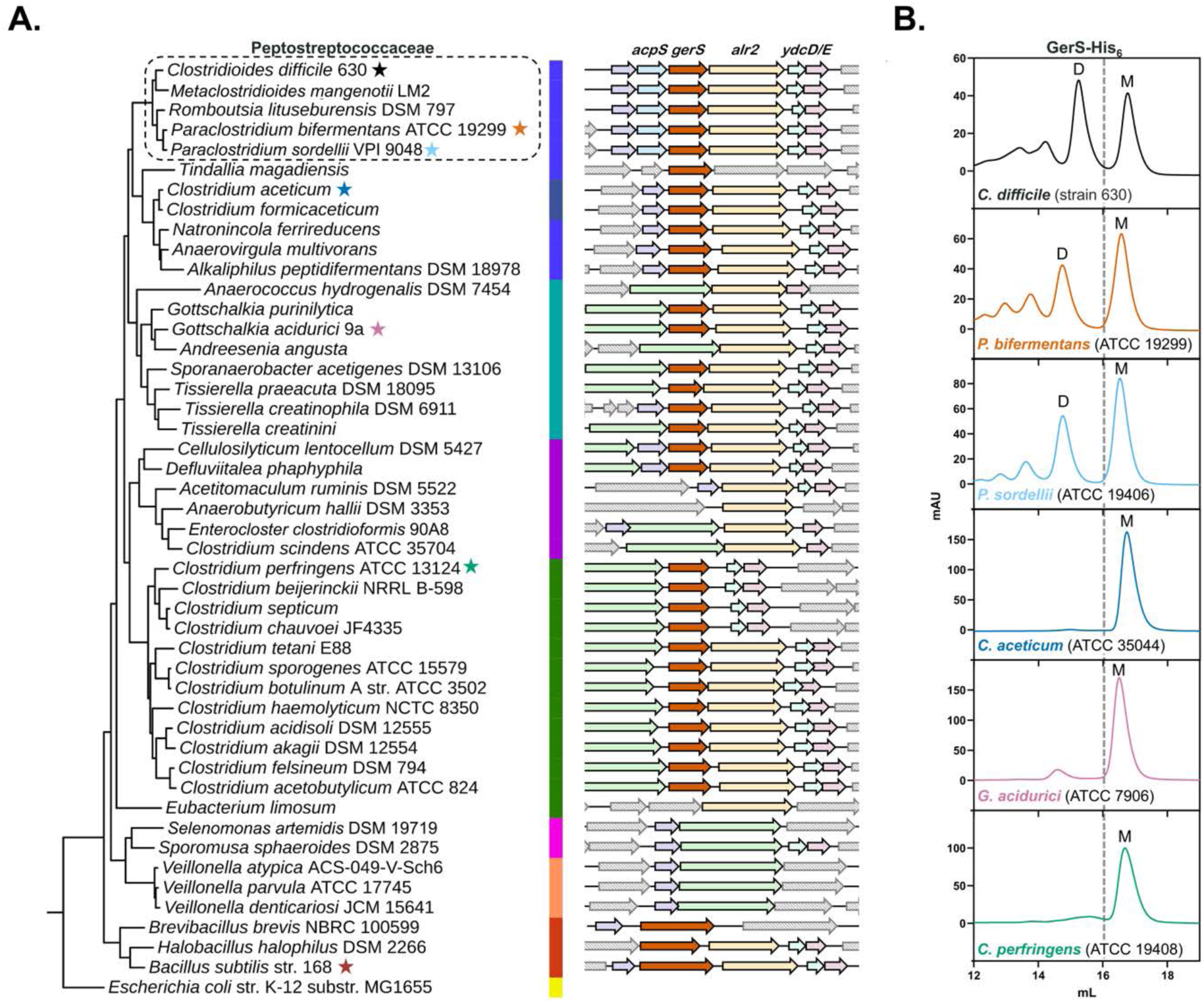
GerS dimerization is conserved in the Peptostreptococcaceae family despite being distributed throughout the Bacilliota phylum. A) Synteny analysis of the conserved *gerS* locus across the orders of the Bacilliota phylum. Peptostreptococcales (blue), Tissierelliales (turquoise), Lachnospirales (purple), Eubacterales (green), Selenomonadales (pink), Vellionellales (orange), Bacillales (red), and the outgroup Enterbacterales (yellow) classes are shown. Colored stars correspond to representative GerS orthologs selected for SEC analysis. B) Size exclusion chromatography (SEC) analysis of GerS orthologs tagged with His_6_ for purification. Peaks are labeled with “D” for dimer or “M” for monomer.

Six orthologs were selected for further characterization from Firmicutes species that sporulate and encode a GerS ortholog. To investigate dimerization, each ortholog was expressed in *E. coli* and purified for analysis via size-exclusion chromatography (SEC). The SEC trace of GerS from *Clostridioides difficile* revealed two distinct peaks, which represent the monomeric and homodimeric forms of the protein ([23], Figure 2B; Supplemental Figure 4).

Notably, two other Firmicutes, *Paraclostridium bifermentans* and *Paraclostridium sordellii* (formerly *Paeniclostridium sordellii* [54]), exhibited similar SEC profiles, indicating that they also form homodimers. In contrast, GerS orthologs from *Clostridium aceticum*, *Gottschalkia acidurici*, and *Clostridium perfringens* displayed a single peak corresponding to the monomeric form. Since *P. bifermentans* and *P. sordellii* are members of the Peptostreptococcaceae family like *C. difficile*, unlike the other species tested, these data suggest that GerS dimerization is specific to the Peptostreptococcaceae family.

We also identified GerS orthologs in the Bacilli class, but they are distinguished by an additional domain of unknown function (DUF4367). We purified the *Bacillus subtilis* ortholog, previously published as SsdC [13], with and without the DUF4367 domain and analyzed the two variants via SEC. Although both traces had two peaks, Coomassie staining of earlier eluting fractions revealed proteins larger than the expected size for SsdC that are likely chaperones (Supplemental Figure 4B-C), indicating that SsdC primarily exists as a monomer *in vitro* and that the DUF4367 domain does not prevent SsdC dimerization. Thus, GerS dimerization is limited to the Peptostreptococcaceae family.

### Mutations in the strand-swap hinge alter GerS dimer stability

To determine the role of the strand swap in regulating GerS dimerization and function, we aimed to identify mutations that destabilize the GerS dimer. Previous studies have shown that mutations that disrupt the ‘hinge’ region of the swapped strand can prevent dimerization (Figure 3A) [28,29,55–57]. While these studies have also shown that it can be difficult to predict which residue will alter a given dimer:monomer equilibrium, we reasoned that replacing the hinge of GerS with the equivalent sequence found in a monomeric PLLLF protein like LolA might prevent GerS dimerization. To this end, we used AlphaFold3 to predict the monomeric structure of GerS and aligned it with the LolA crystal structure (PDB 1IWL). This alignment revealed that the GerS strand-swap hinge, Lys71-Thr72-Asp73-Asn74 (KTDN), lies in a similar position to LolA Arg64-Pro65-Asn66-Leu67 (RPNL; Figure 3B). We then tested the effect of replacing individual amino acids in the *C. difficile* GerS hinge with the analogous residues in the *E. coli* LolA region. Intriguingly, replacing Thr72 with Pro increased the proportion of GerS in the monomeric form, while replacing Lys71 with Arg and Asn74 with Leu increased the dimeric form (Figure 3C; Supplemental Figure 5A). In contrast, the GerS^D73N^ variant precipitated quickly after purification and co-purified with the chaperones from *E. coli* (Supplemental Figure 5B)[58–60], strongly suggesting that the substitution destabilized GerS.

**Figure 3.**
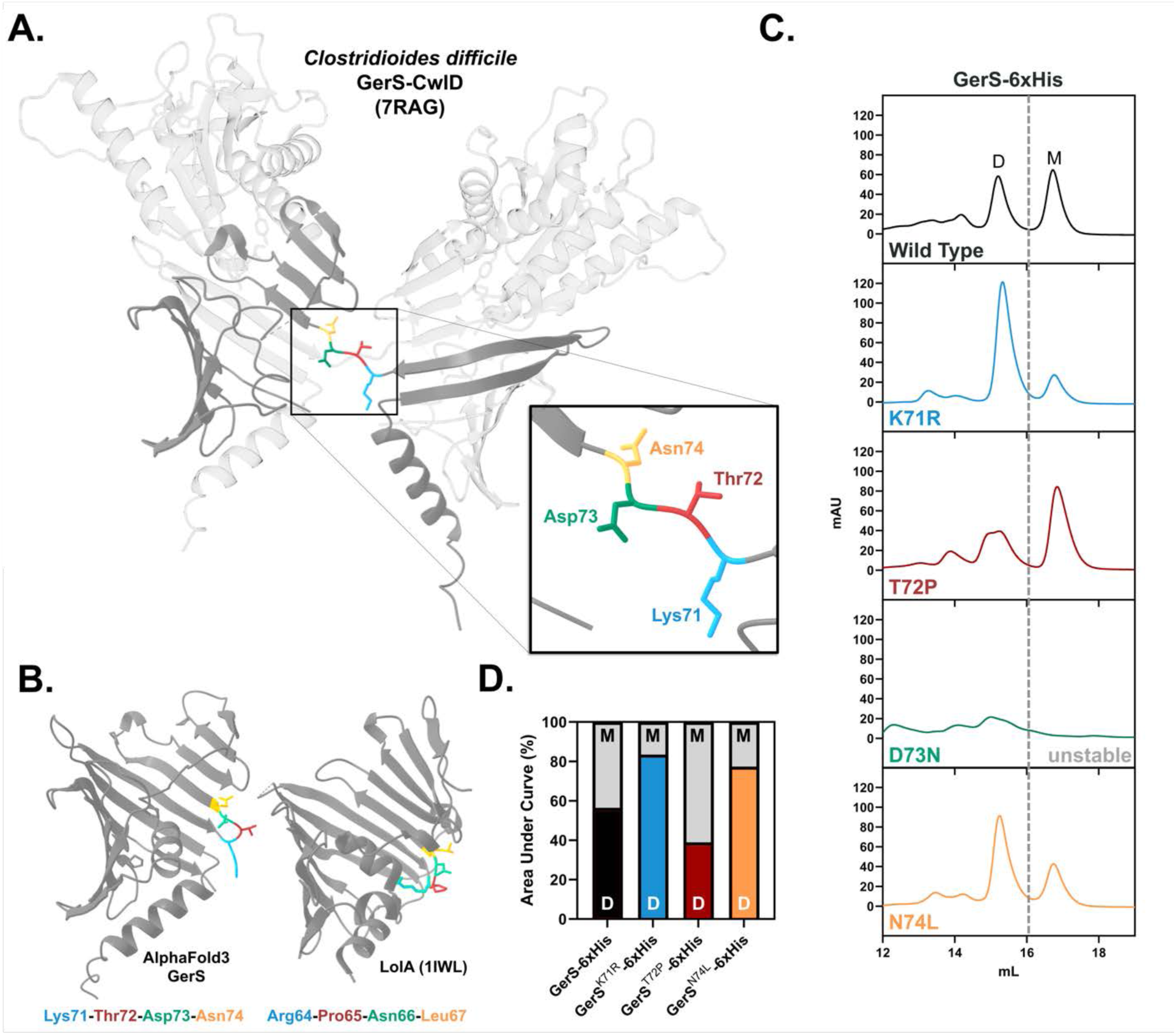
GerS hinge mutations guided by the LolA structure modulate its dimer:monomer equilibrium. A) Ribbon representation of the crystal structure of the GerS-CwlD complex (PDB 7RAG). The hinge is colored by the residue shown in the inset. B) Ribbon representation of the AlphaFold3 GerS structure and LolA crystal structure (PDB - 1IWL). Hinge of GerS and analogous residues on LolA are colored by aligned residues. C) Size exclusion chromatography (SEC) traces of each GerS variant tagged with His_6_. Peaks are labeled with “D” for dimer or “M” for monomer. D) Ratio of the areas under each peak as a percentage of the total area measured. “D” is the area under the dimer peak, and “M” is the area under the monomer peak.

Since the GerS^K71R^ and GerS^N74L^ variants favored the dimeric form relative to WT GerS (Figure 3C-D), we purified the dimeric fractions and re-analyzed them by SEC to compare the relative stabilities of the GerS homodimers formed by each variant. The re-purified GerS^K71R^ or GerS^N74L^ variants again showed a higher dimer-to-monomer ratio than WT GerS (Supplemental Figure 6A-B), indicating that the K71R and N74L substitutions enhance the stability of the GerS homodimer structure.

### Proline substitutions in the strand-swap hinge destabilize the GerS homodimer

Since GerS^T72P^ favors the monomeric form compared to WT GerS (Figure 3C-D), we next tested the effect of changing the other hinge residues to proline. Previous studies have found that introducing prolines into the strand-swap hinge affects dimerization in a wide range of proteins across all domains of life [28,29]. For example, highly conserved prolines in the loops of *Mycobacterium tuberculosis* BlaC maintain BlaC in a monomeric and functional form, whereas substitutions of these residues result in dimerization and loss of function [61]. When the other hinge residues of GerS were replaced with proline (K71P, D73P, and N74P), all three substitutions appeared to destabilize GerS (Figure 4A), and the resulting variants were quick to crash out of solution. The D73P and N74P substitutions resulted in low protein yields, while SEC analyses of the GerS^K71P^ variant revealed that it copurified with *E. coli* proteins likely to be chaperones (Supplemental Figure 7B). Thus, the location of the proline substitution is critical for regulating the GerS dimer-to-monomer equilibrium.

**Figure 4.**
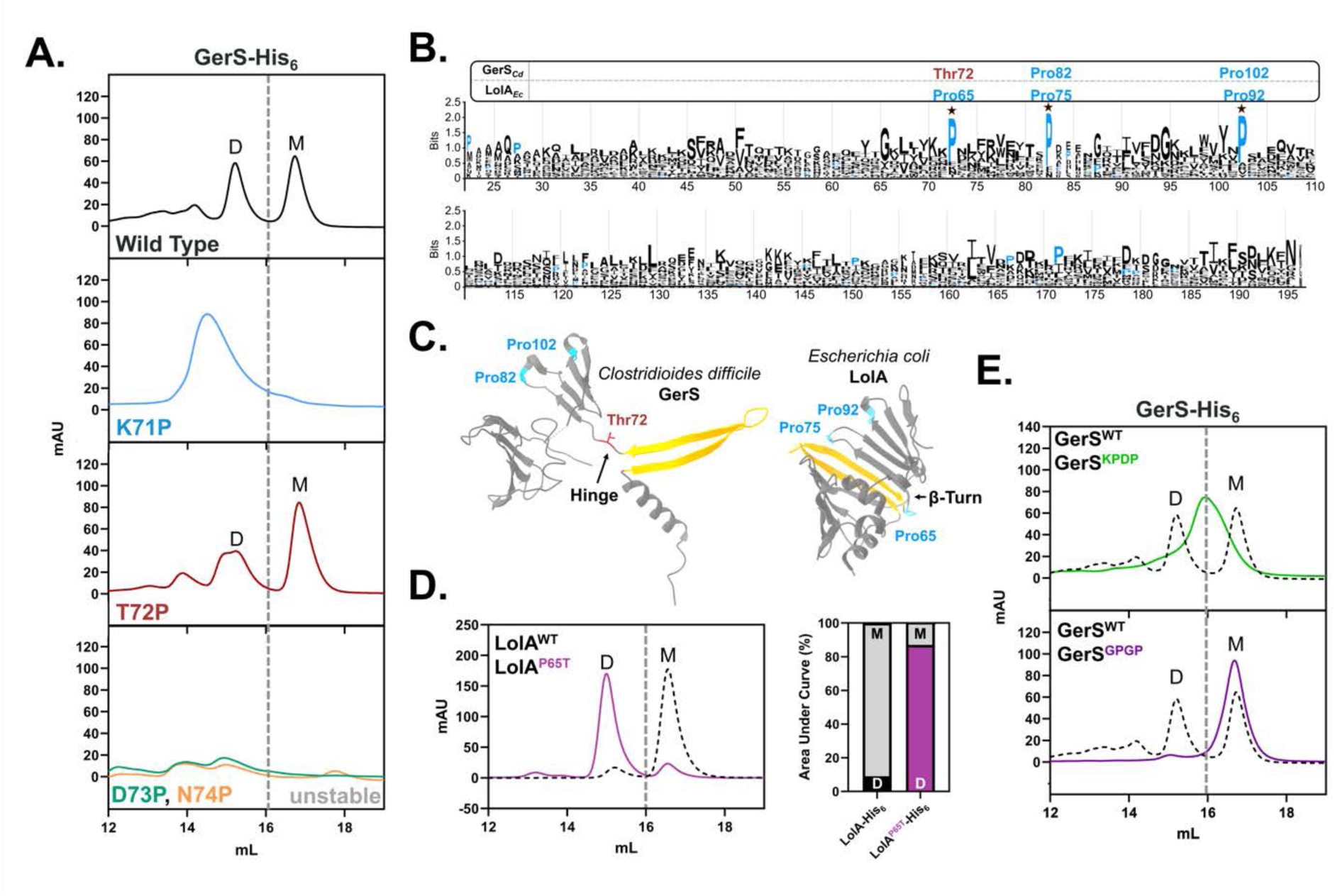
Prolines in the hinge region of PLLLF proteins reduce dimerization. A) Size exclusion chromatography (SEC) traces of His_6_-tagged GerS variants carrying single proline substitutions in the hinge region. B) Sequence logo depicting the frequency of amino acids found at a given position based on DALI structural alignments of 17 representative PLLLF proteins [48]. Prolines are highlighted in blue. (C) Amino acids aligned to each conserved proline in GerS and LolA are displayed in the ribbon representations of the crystal structures shown (PDB 7RAG and 1IWL, respectively). D) SEC trace of LolA^WT^ and LolA^P65T^ with area under the curve calculations. E) SEC trace of dual proline GerS variants. Peaks are labeled with either “D” for dimer or “M” for monomer.

Consistent with this finding, analysis of the conservation of hinge prolines among PLLLF proteins using the DALI alignment revealed that three proline residues are highly conserved among this superfamily, and each of these prolines correspond to beta-turns in PLLLF proteins (Figure 4B). Notably, GerS contains two of these conserved prolines but not the conserved proline at residue 72 (Figure 4C). Together, these observations highlight how a proline at this location destabilizes the GerS homodimer and promotes GerS monomer formation.

To test whether the conserved proline in the analogous position of other monomeric PLLLF proteins regulates dimerization, we substituted the *E. coli* LolA Pro65 with a Thr (P65T). Remarkably, while wild-type LolA eluted as a monomer, LolA^P65T^ eluted primarily as a dimer (Figure 4D). Taken together, these data highlight how a proline at this site promotes β-turn formation and thus the monomeric form, consistent with past studies of β-turns [62,63].

While the T72P substitution effectively shifted the dimer-to-monomer equilibrium toward the monomer state, it did not completely prevent the formation of GerS homodimers (Fig 3C, 4A). To further destabilize the dimer, we introduced a second proline at the fourth residue, creating the hinge sequence Lys71-Pro72-Asp73-Pro74 (KPDP). Since prolines in β-turns are often paired with flexible glycine residues to counteract their rigidity [63–65], we also replaced the first and third residues in the hinge with glycines, yielding a Gly71-Pro72-Gly73-Pro74 (GPGP) hinge sequence. The GerS^KPDP^ variant eluted as a single peak at a volume that falls between the dimer and monomer peaks, suggesting that the variant interconverts between the monomer and dimer forms as it is resolved on the column, whereas the GerS^GPGP^ variant exclusively eluted as a monomer (Figure 4E). These results indicate that the addition of a second proline in the strand-swap hinge further destabilizes the GerS homodimer—favoring the monomeric form—especially when glycines are introduced to provide greater conformation flexibility.

### Decreasing GerS dimer formation reduces its affinity for CwlD and impairs *C. difficile* spore germination

Since we originally hypothesized that strand-swapping in GerS enhances its ability to bind CwlD based on analyses of other strand-swapped proteins [42], we next tested if destabilizing the homodimer affects its affinity for CwlD. To this end, we co-affinity purified His_6_-tagged CwlD with untagged GerS variants and analyzed their interactions by SEC. Untagged GerS^WT^, GerS^K71R^, and GerS^N74L^ co-purified efficiently with His_6_-tagged CwlD and eluted as the GerS:CwlD complex as well as the GerS and CwlD monomers in SEC analyses (Figures 5A,B, Supplementary Figure 8). It should be noted that the GerS:CwlD complex eluted as several peaks due to the higher resolution SEC performed relative to our prior work [23].

**Figure 5.**
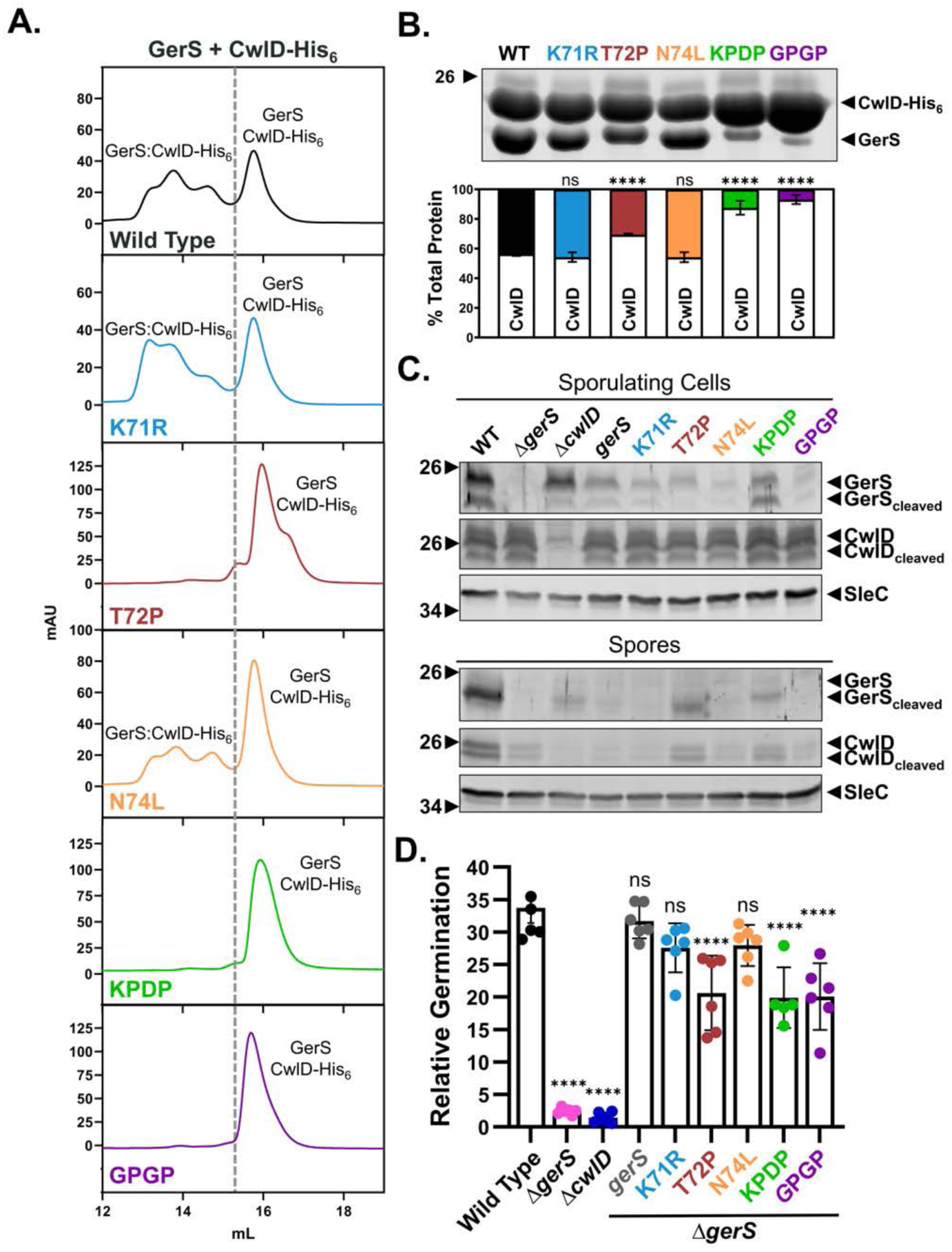
GerS substitutions that reduce dimerization decrease binding to CwlD and spore germination. A) SEC traces of GerS hinge region variants copurified with His_6_-tagged CwlD. Peaks labeled with “GerS:CwlD-His_6_”, correspond to the complex formed between GerS and CwlD-His_6_. Peaks labeled with “GerS” and “CwlD-His_6_”, represent the individual proteins that elute outside of complex. B) Coomassie stain of untagged GerS variants copurified with His_6_-tagged CwlD. C) Western blot analyses of sporulating cell or purified spore lysates prepared from Δ*gerS* complemented with constructs encoding GerS hinge region variants. SleC was used as a loading control. D) Relative germination based on the change in optical density (OD_600_) of purified spores over time following the addition of 1% (19 mM) taurocholate in rich media. Germination in arbitrary units was calculated using the area below inverted OD_600_ curves. Each *gerS* mutant is expressed by its native promoter in the *pyrE* locus in a Δ*gerS* background. A one-way ANOVA followed by Dunnett’s multiple comparisons test was used to compare the mean of each *gerS* mutant to the Δ*gerS*/*gerS* complement strain based on a minimum of three biological replicates. Significance relative to WT is shown. (**** p < 0.00005).

Since the K71R and N74L substitutions stabilized GerS homodimerization, we considered the possibility that these variants may bind CwlD more tightly than wild-type GerS. To test this, we resolved co-purifications of GerS^K71R^:CwlD-His_6_ and GerS^N74L^:CwlD-His_6_ using SEC, collected the dimer fractions, and re-resolved them through SEC again (Supplemental Figure 9). To our surprise, the GerS^K71R^ and GerS^N74L^ variants formed weaker interactions with CwlD-His_6_ than wild-type GerS. However, the differences between GerS^K71R^ and GerS^WT^ were subtle. We next attempted to further stabilize the GerS dimer by replacing the GerS hinge with the hinge (QVVAG) from the strand-swapping cystatin, Stefin B, because this hinge was previously reported to reliably form domain-swapped homodimers [66]. However, substituting GerS’s hinge region for Stefin B’s hinge region (GerS^QVVAG^) destabilized GerS, causing it to quickly crash out of solution; indeed, the SEC trace matched that of the unstable D73N substitution (Supplemental Figure 10), and untagged GerS^QVVAG^ failed to copurify with CwlD-His_6_ (Supplemental Figure 10B). Therefore, the effect of stabilizing the GerS homodimer on CwlD remains unclear.

In contrast to GerS^K71R^ and GerS^N74L^, considerably less GerS^T72P^ co-purified with His_6_-tagged CwlD (Figure 5A, Supplementary Figure 7). In addition, when the co-purification was resolved by SEC, no GerS^T72P^:CwlD-His_6_ complex was observed, indicating that the co-affinity purified complex was not stable during SEC (Supplementary Figure 8). These latter data indicate that destabilizing the GerS homodimer decreases its binding to CwlD. Consistent with this conclusion, the GerS^KPDP^ and GerS^GPGP^ variants, which favor the monomeric form (Figure 4E), had even lower affinity for CwlD-His_6_ (Figure 5B).

To determine the impact of preventing GerS dimerization on GerS function during *C. difficile* germination, we complemented our Δ*gerS* mutant with constructs encoding dimer-breaking GerS mutations and measured their effects on GerS stability and spore germination efficiency. While GerS^K71R^ and GerS^N74L^ interacted with CwlD at the same levels as wild-type GerS *in vitro*, both variants were detected at lower levels in sporulating cells compared to the Δ*gerS* wild-type complement. Additionally, both mutants were almost undetectable in purified spores (Figure 5C). Consistent with our previous findings that interaction with CwlD stabilizes GerS [23], very low levels of the GerS^GPGP^ variant were detected in sporulating cells, and we were unable to detect this variant in purified spores (Figure 5C). However, the GerS^T72P^ variant and GerS^KPDP^ variant were readily detectable in both sporulating cells and purified spores, implying that their ability to interact with CwlD is sufficient to stabilize the GerS variants in spores (Figure 5C).

Notably, even though the dimer-disrupting strand-swap mutations decreased GerS levels in mature spores, the GerS dimerization mutants exhibited only an ∼2-fold decrease in germination (p < 0.00005, Figure 5D). This finding is consistent with our prior work indicating that a GerS mutation, D106R, which disrupts a salt-bridge at the interaction interface of GerS and CwlD, results in a modest but significant decrease in germination [23]. Our results also suggest that relatively GerS is needed in spores to license CwlD amidase activity.

### Interaction with CwlD requires both strand swapping and a salt bridge

Since our analyses revealed that GerS mutants defective in dimerization still allow for *C. difficile* germination to proceed, additional features of GerS beyond strand-swapping allow GerS to bind CwlD. Indeed, our *in vitro* analyses revealed that GerS dimerization mutants can still co-purify with CwlD-His_6_, albeit at greatly reduced levels (Figure 5B). Since we previously showed that the interaction between GerS and CwlD is partially dependent on a salt bridge that forms between arginine 169 of CwlD and aspartate 106 of GerS [23], we reasoned that GerS binding to CwlD may depend on both the salt bridge and GerS strand swap (Figure 6A). To test this possibility, we combined the salt-bridge and strand-swapping mutations to generate GerS^T72P,D106R^, GerS^KPDP,D106R^, and GerS^GPGP,D106R^ variants. Remarkably, these combined mutations fully disrupted GerS binding to CwlD, since the untagged GerS variants were not detectable in the co-elutions with His_6_-tagged CwlD (Figure 6B). Consistent with this finding, purified spores from Δ*gerS* strains complemented with constructs encoding the combined salt-bridge and strand-swap mutations failed to germinate (Figure 6C), presumably due to their inability to bind CwlD and allosterically activate its amidase activity during cortex modification [23]. In addition, the combined mutations greatly decreased GerS levels even in sporulating cells (Figure 6D). Taken together, our data highlight the importance of both the strand swap and salt bridge on GerS binding to CwlD and, by extension, regulation of *C. difficile* spore germination.

**Figure 6.**
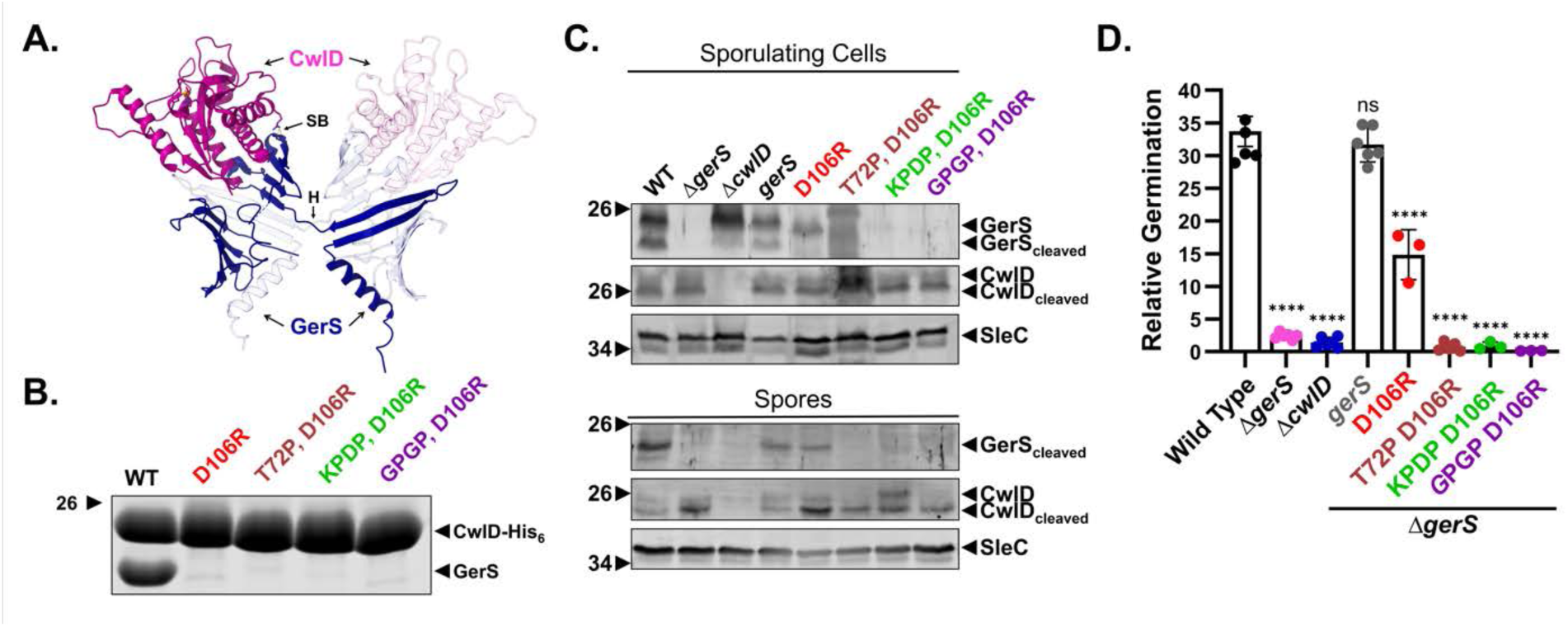
GerS interaction with CwlD is dependent on both strand-swapping and a salt bridge. A) GerS:CwlD structure (7RAG). SB, Salt bridge; H, Hinge. B) Coomassie-stained eluates of untagged homodimer-disrupting GerS variants combined with the salt bridge mutation D106R copurified with His_6_-tagged CwlD. C) Relative germination calculated by area under the percent change in OD600 curve of spores in BHIS in the presence of 1% taurocholate over 1.5 hours. Each *gerS* mutant is expressed by its native promoter in the *pyrE* locus in a Δ*gerS* background. A one-way ANOVA followed by Dunnett’s multiple comparisons test was used to compare each *gerS* mutant mean to the mean Δ*gerS*/*gerS* complement strain. Each mean is calculated from at least three biological replicates. Germination assay and statistics were run with Figure 5D, therefore Wild Type, Δ*gerS*, Δ*cwlD,* and Δ*gerS/gerS* complement are the same between both panels. The significance relative to WT is shown. (**** p < 0.00005). D) Detection of GerS variants by western blotting of sporulating cell or purified spore lysates. Polyclonal antibodies against GerS, CwlD, and SleC were used.

## Discussion

Our prior discovery that the *C. difficile* GerS lipoprotein binds the CwlD amidase as a strand-swapped dimer led to the question of whether strand-swapping by GerS enhances its ability to allosterically activate the CwlD amidase [23]. By identifying mutations that shift the GerS dimer:monomer equilibrium to the monomeric form (Figure 3), we show that monomeric GerS has greatly reduced affinity for its binding partner, CwlD, which in turn reduces, but does not abolish, *C. difficile* spore germination (Figure 5). Since combining monomer-promoting mutations with a mutation that disrupts a salt-bridge at the GerS:CwlD interface abrogates GerS’s ability to activate the CwlD amidase, our data reveal a critical role for GerS’s hinge region in regulating the allosteric activation of CwlD.

While our findings are consistent with prior work highlighting that strand-swapping can stabilize adhesive interactions, such as in cadherins and the LolA^F68E^ variant, they also raise the possibility that the dynamic instability of the strand-swapped dimer, which exists in near-equal proportions with the monomeric form *in vitro* (Figure 3), contributes to GerS function. Strand swapping is sometimes the result of localized unfolding around the hinge. While the rest of the protein folds, the strand to be swapped unfolds, leading to monomeric and dimeric conformations [32,67]. Like GerS, the yeast cell-cycle regulation protein p13^suc1^ has both a native monomer and dimer state [29]. Two prolines in the hinge generate tension in each state, functioning as a sort of spring between both the monomer and dimer conformation.

Substitutions in the hinge of the swapped structure shift the equilibrium of the dimer and monomer populations [29]; in a similar manner, the K71R and N74L substitutions in GerS stabilize the dimeric conformation, while the T72P and GPGP substitutions stabilize the monomeric conformation. The persistence of the two stable populations in near equal proportions suggests that the GerS monomer may be functionally important, perhaps to ensure that GerS transiently interacts with CwlD during MAL formation. Alternatively, additional regulatory factors may influence or stabilize one conformation over the other during this process.

Our finding that replacing the GerS strand-swap hinge with the corresponding region from Stefin B [66] destabilizes the protein (Supplemental Figure 10) suggests that dimerization in these proteins depends on distinct, non-interchangeable hinge features. While most domain-swapped proteins have hinge loops of four to five residues, they otherwise vary widely in sequence and structure. Nonetheless, a common feature is that these hinges originate from β-turns in their monomeric structures [38,68,69]. Consistent with previous findings that conserved prolines in β-turns can prevent deleterious strand-swapping in *M. tuberculosis* BlaC [61], we found that the introduction of prolines into the GerS hinge destabilizes the strand-swapped dimer (T72P, GPGP; Figure 4)[61]. Yet, there are also examples where introducing prolines into β-turns instead promotes domain swapping [70–72]. Together, these findings underscore that there is no universal “syntax” for strand swapping and that structural states must be empirically determined on a case-by-case basis.

The fact that a single threonine-to-proline substitution in the GerS hinge dramatically alters its conformational stability implies that PLLLF proteins may have evolved inherent instability at this site, potentially enabling their structural flexibility. Although *C. difficile* and *E. coli* diverged billions of years ago [73], a reciprocal substitution at the same hinge position in LolA (LolA^P65T^) is sufficient to shift the protein from a monomer to a strand-swapped dimer (Figure 4). This evidence, along with the similarly strand-swapped LolA^F68E^ (published F47E) [41,42] structure, supports the idea that the LolA monomeric structure around these sites is intrinsically unstable. These findings also highlight the ease with which PLLLF proteins can evolve strand-swapped dimerization. This finding, along with the grouping of both proteins into the same superfamily by SCOP [74], also raises the intriguing possibility of a shared ancestry between LolA and GerS. During the evolution of monoderm bacteria, which involved the loss of the ancestral outer membrane [75–77], there would no longer be a need to transport lipoproteins across the periplasmic space, facilitating a specialization of the ancestral proteins during sporulation in endospore-forming monoderms [13,23]. This hypothesis may also clarify why PLLLF proteins are absent in certain non-spore-forming monoderms such as *Staphylococcus aureus* and the seemingly convergent roles of LppX [10] and Lpr proteins [11,12] in transporting glycolipids across the mycobacterial periplasm.

Much work remains to be done to understand precisely why GerS strand swapping stabilizes its interaction with CwlD. Notably, CwlD binding to its Zn^2+^ co-factor also stabilizes the GerS:CwlD interaction, so it is possible that Zn^2+^-bound CwlD adopts a conformation that similarly promotes GerS strand-swapping. Identifying the allosteric circuit that allows GerS binding to CwlD to be communicated to CwlD’s active site and vice versa will likely provide insight into the role of GerS strand-swapping in the allosteric activation of CwlD. Another question that arises from our identification of GerS orthologs in other clostridial species that do not form dimers is whether these GerS orthologs have additional or different conserved functions beyond regulating CwlD activity during cortex modification? The requirement of CwlD for GerS processing and incorporation into the spore may indicate that GerS functions in a very specialized way in the Peptostreptococcaceae family [22]. In contrast, it is possible that GerS orthologs in other Clostridia function similarly to SsdC in *B. subtilis*, which is required for proper spore morphology through its impacts on spore cortex assembly [13]. Nevertheless, how these orthologs are incorporated into the spore if they do not interact with CwlD as well as whether GerS and CwlD can independently localize to the forespore remain open questions and will require analyses in organisms other than *C. difficile*.

## Materials and Methods

### Structural alignment, sequence analysis, and phylogenetics

17 PLLLF proteins were selected from the InterPro [44] SuperFamily SSF89392, representing the taxonomic diversity of the PLLLF protein family. AlphaFold3 [47] was used to predict the structure of each protein. The 17 proteins were aligned using DALI [48], which generates an amino-acid sequence alignment based on the closest amino acids in each aligned structure. The structural alignment was imported into Geneious Prime 2024.0.5. to calculate the %identity and %similarity between each sequence (see Table S1). For %similarity calculations, the Blosum45 matrix was used with a threshold of 0. WebLogo [78] was used to generate the sequence logo of the structural alignment.

The phylogenetic tree of the major bacterial classes, with Archaea and Eukarya as used as outgroups, was constructed using genomes publicly available on KBase [79]. Five genomes for each class were selected, although there were several instances when five were not available for a given class. The complete list of these genomes can be found in Table S2. The KBase app “Insert Genome into SpeciesTree - v2.2.0” was used to identify, concatenate, and align a set of 49 core, universal genes defined by COG (Clusters of Orthologous Groups) gene families across all selected genomes. The app generates the phylogenetic tree using FastTree2 [80]. The tree was visualized and prepared into a figure using iTOL [81]. Identification of PLLLF proteins in each class was done by searching for PLLLF proteins encoded by members of each class in the InterPro [44] SuperFamily SSF89392 and by running FoldSeek [43] using the LolA crystal structure (1IWL).

### Bacterial strains and growth conditions

*C. difficile* strains 630Δ*ermΔcwlD*Δ*pyrE* and 630Δ*ermΔgerS*Δ*pyrE* were used as the parental strains for *pyrE*-based allelic-coupled exchange (ACE) [82], which allows for single-copy complementation from the ectopic *pyrE* locus. The *C. difficile* strains used are listed in Table S3. They were grown on brain heart infusion media (BHIS) supplemented with taurocholate (TA, 0.1% wt/vol; 1.9 mM), thiamphenicol (10-15 µg /mL), kanamycin (50 µg/ml), cefoxitin (8 µg/ml) and L-cysteine (0.1% w/v; 8.25 mM) as needed. For ACE, *C. difficile* was grown on *C. difficile* defined medium (CDDM) [83]. Cultures were grown at 37°C under anaerobic conditions using a gas mixture containing 85% N_2_, 5% CO_2_ and 10% H_2_.

*Escherichia coli* strains used for BL21(DE3)-based protein production and HB101/pRK24-based conjugations are listed in Table S3. *E. coli* strains were grown at 37°C with shaking at 225 rpm in Luria-Bertani broth (LB). The media was supplemented with chloramphenicol (20 µg/ml), ampicillin (50 µg/ml), or kanamycin (30 µg/ml) as indicated.

### *E. coli* strain construction

Plasmid constructs were confirmed by sequencing using Genewiz or Plasmidsaurus (see the Benchling links in Table S3) and transformed into either the HB101/pRK24 conjugation strain (used with *C. difficile*) or BL21(DE3) expression strain.

### *C. difficile* strain construction and complementation

Allele-couple exchange (ACE) was used to construct all complementation strains as previously described by conjugating HB101/ pRK24 carrying pMTL-YN1C plasmids into Δ*pyrE-*based strains.

### Protein purification

Starter cultures were grown in 20 mL LB broth with 30 µg/mL of kanamycin and, in some cases, 100 μg/mL ampicillin. Terrific broth (TB) with ampicillin and kanamycin was inoculated with the starter culture (1:1000) and incubated for ∼60 hr at 20°C with 225 rpm shaking. The purifications were carried out in two steps: nickel bead-based (co-)affinity purification followed by size exclusion chromatography (SEC). Affinity purification was carried out as stated above: the cells were pelleted, resuspended in lysis buffer (500 mM NaCl, 50 mM Tris [pH 7.5], 15 mM imidazole, 10% [vol/vol] glycerol, 2 mM ß-mercaptoethanol), flash frozen in liquid nitrogen, thawed and then sonicated. The insoluble material was pelleted, and the soluble fraction was incubated with Ni-NTA agarose beads (Qiagen) for 2 hours to capture the His_6_-tagged proteins. Washes were carried out five times with low imidazole buffer (500mM NaCl, 10mM Tris-HCl pH 7.5, 10% (v/v) glycerol, 15 mM imidazole, 2 mM ß -mercaptoethanol) to decrease non-specific binding to the beads. Proteins were eluted using high-imidazole buffer (500 mM NaCl, 50 mM Tris [pH 7.5], 200 mM imidazole, 10% [vol/vol] glycerol, 2 mM ß-mercaptoethanol) after nutating the sample for 5 to 10 min. Pooled protein elutions were concentrated to 20 mg/ml or less in a size exclusion chromatography buffer consisting of 200 mM NaCl, 10 mM Tris HCl pH 7.5, 5% glycerol, and 1 mM DTT.

### Size exclusion chromatography (SEC)

SEC was carried out using a Superdex 200 Increase 10/300 GL (GE Healthcare) column. A high-resolution protocol was used to resolve affinity-purified protein elutions by SEC. Specifically, purified proteins were diluted to a final concentration of 10 mg/mL in the SEC buffer listed above, and then 100 µL of the 10 mg/mL diluted protein was resolved using a 0.2 mL/min flow rate. The resulting fractions were resolved using SDS-PAGE, and the gels were Coomassie stained using GelCode Blue according to the manufacturer’s instructions (ThermoFisher Scientific). Protein sequencing of non-GerS bands was performed by automated Edman degradation on an ABI Procise 494 HT Protein/Peptide Sequencer at the Tufts University Core Facility (TUCF).

### Western blot analysis

Samples for immunoblotting were prepared by resuspending sporulating cell pellets in 100 µl of PBS, and freeze-thawing a 50 µL aliquot of this resuspension for three cycles. The pellet was then resuspended in 100 µL EBB buffer (8 M urea, 2 M thiourea, 4% [wt/vol] SDS, 2% [vol/vol] ß-mercaptoethanol). *C. difficile* spores (∼1 x 10^6^) were resuspended in EBB buffer, which extracts proteins in all layers of the spore, including the core. Both sporulating cells and spores were incubated at 95°C for 20 min with vortex mixing. Samples were centrifuged for 5 min at 15,000 rpm, and 4 x sample buffer (40% [vol/vol] glycerol, 1 M Tris [pH 6.8], 20% [vol/vol] ß-mercaptoethanol, 8% [wt/vol] SDS, 0.04% [wt/vol] bromophenol blue) was added. Samples were incubated again at 95°C for 5 to 10 min with vortex mixing followed by centrifugation for 5 min at 15,000 rpm. The samples were resolved by the use of 14% SDS-PAGE gels and transferred to a Millipore Immobilon-FL polyvinylidene difluoride (PVDF) membrane. The membranes were blocked in Odyssey blocking buffer with 0.1% (vol/vol) Tween 20. Polyclonal rabbit anti-GerS and anti-CwlD antibodies were used at a 1:500 and 1:750 dilution, respectively. Polyclonal mouse anti-SleC antibodies were used at a 1:5000 dilution. IRDye 680CW and 800CW infrared dye-conjugated secondary antibodies were used at 1:30,000 dilutions. An Odyssey LiCor CLx was used to visualize secondary antibody infrared fluorescence emissions.

### Spore purification

Sporulation was induced on 70:30 agar plates for 3-5 days, and the cultures were harvested for spore purifications. Spores were washed 6 times in ice-cold water, incubated overnight in water on ice, treated with DNase I (New England Biolabs) at 37°C for 60 min, and purified on a HistoDenz (Sigma-Aldrich) gradient. Phase-contrast microscopy was used to assess spore purity (>95% pure); the optical density of the spore stock was measured at OD_600_; and spores were stored in water at 4°C. Four 70:30 plates were used to induce sporulation for each strain; purified spores were resuspended in 200 µL, and the total optical density of this mixture was determined for each strain. At least three independent spore preparations were performed. Spore purity as assessed by phase-contrast microscopy. Only spore suspensions that were >95% pure were used.

### Optical density-based germination assay

For OD_600_-based germination assays, germination was induced with taurocholate in 96-well plates. Specifically, ∼1.6 x 10^7^ spores (0.55 OD_600_ units) for each condition tested were resuspended in BHIS and then 180 μL of this suspension was aliquoted in duplicate into a 96 well flat-bottom tissue culture plate (Falcon) for each condition tested. The spores were exposed to 1% taurocholate (19 mM), or water (untreated) in a final volume of 200 μL. The OD_600_ of the sample was measured every 3 minutes in a Synergy H1 microplate reader (Biotek) at 37 °C with constant shaking between readings. The change in OD_600_ over time was calculated as the ratio of the OD_600_ at each time point to the OD_600_ at time zero. Relative germination was calculated by measuring the area between the drop in OD_600_ and 1.0 after 39 minutes (13 reads). These assays were performed on 3-6 independent spore preparations depending on the condition. Data shown are averages from these replicates. Comparison of each mean to WT was performed using a one-way ANOVA followed by Dunnett’s multiple comparisons test, with a single-pooled variance.

