## Supplemental figures and tables for "Strand swapping in the LolA-like protein GerS promotes the allosteric activation of a bacterial amidase in *Clostridioides difficile*"

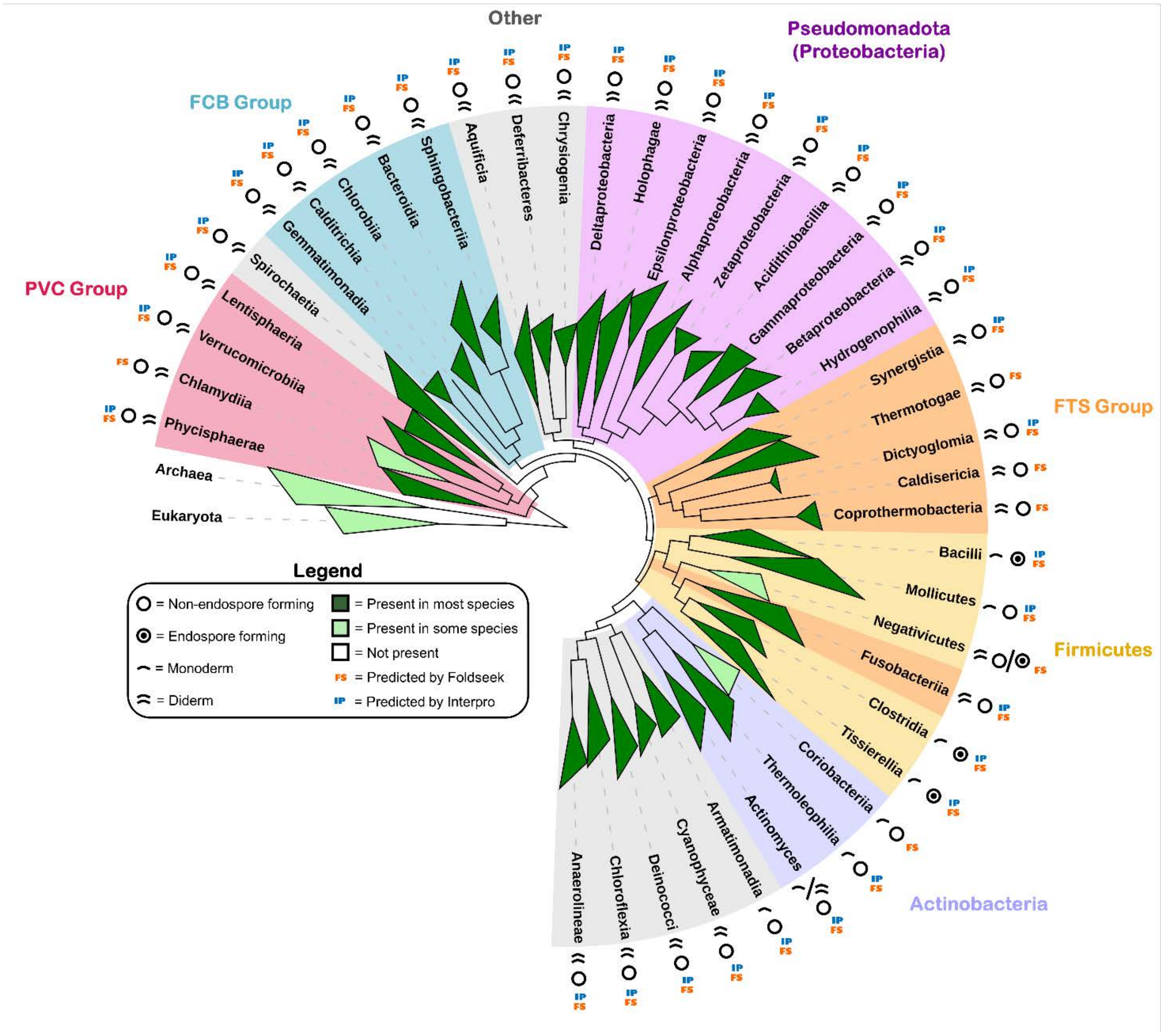


**Supplemental Figure 1. Presence of PLLLF proteins is widespread throughout major bacterial classes.** FastTree [80] alignment of 51 concatenated core genes from five representatives (when available) in each major bacterial class. Each class is labeled as either being made up of monoderms or diderms as well as endospore forming or non-endospore forming. Both FoldSeek [43] and InterPro [44] were used to find PLLLF proteins in each class. Clades are shaded either light green or dark green depending on whether a few or most of the members encode PLLLF proteins.

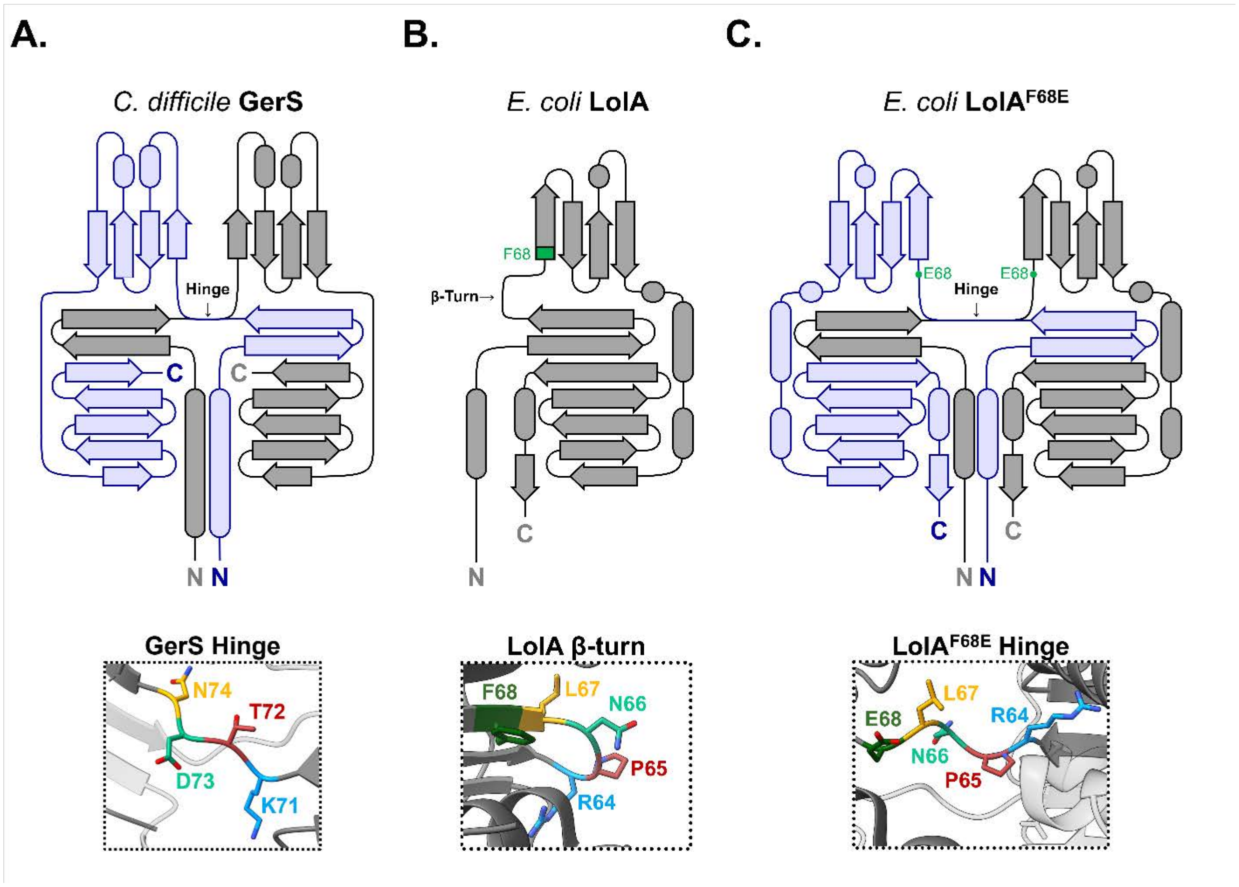


**Supplemental Figure 2. Two-dimensional topology diagrams of GerS and LolA$^{F68E}$.** Topology diagrams depicting secondary structure of GerS (A, PDB 7RAG) and the *Escherichia coli* LolA variants LolA$^{WT}$ (B, PDB 1IWL) and LolA$^{F68E}$ (C, PDB 6FHM). The diagrams were modified from PDBsum [85]. Insets show the hinge or β-turn from crystal structures corresponding to each topology diagram.

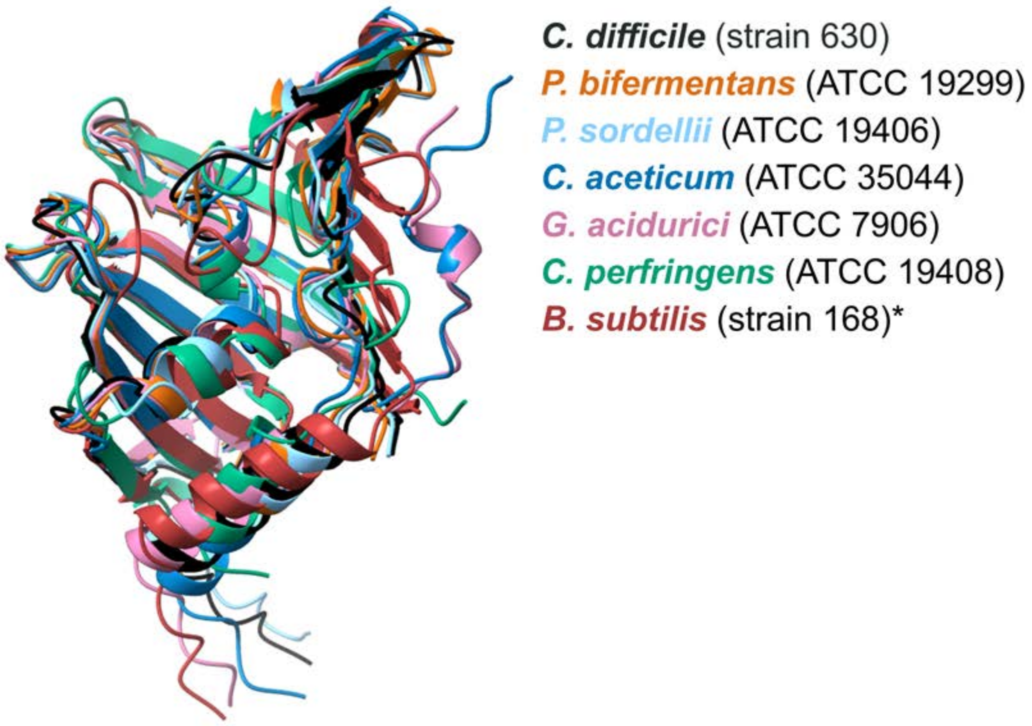


**Supplemental Figure 3. Alignment of predicted structures of GerS orthologs.** AlphaFold3-predicted structures of the seven orthologs analyzed using size exclusion chromatography. SsdC (maroon) is aligned here without the DUF4367 domain.

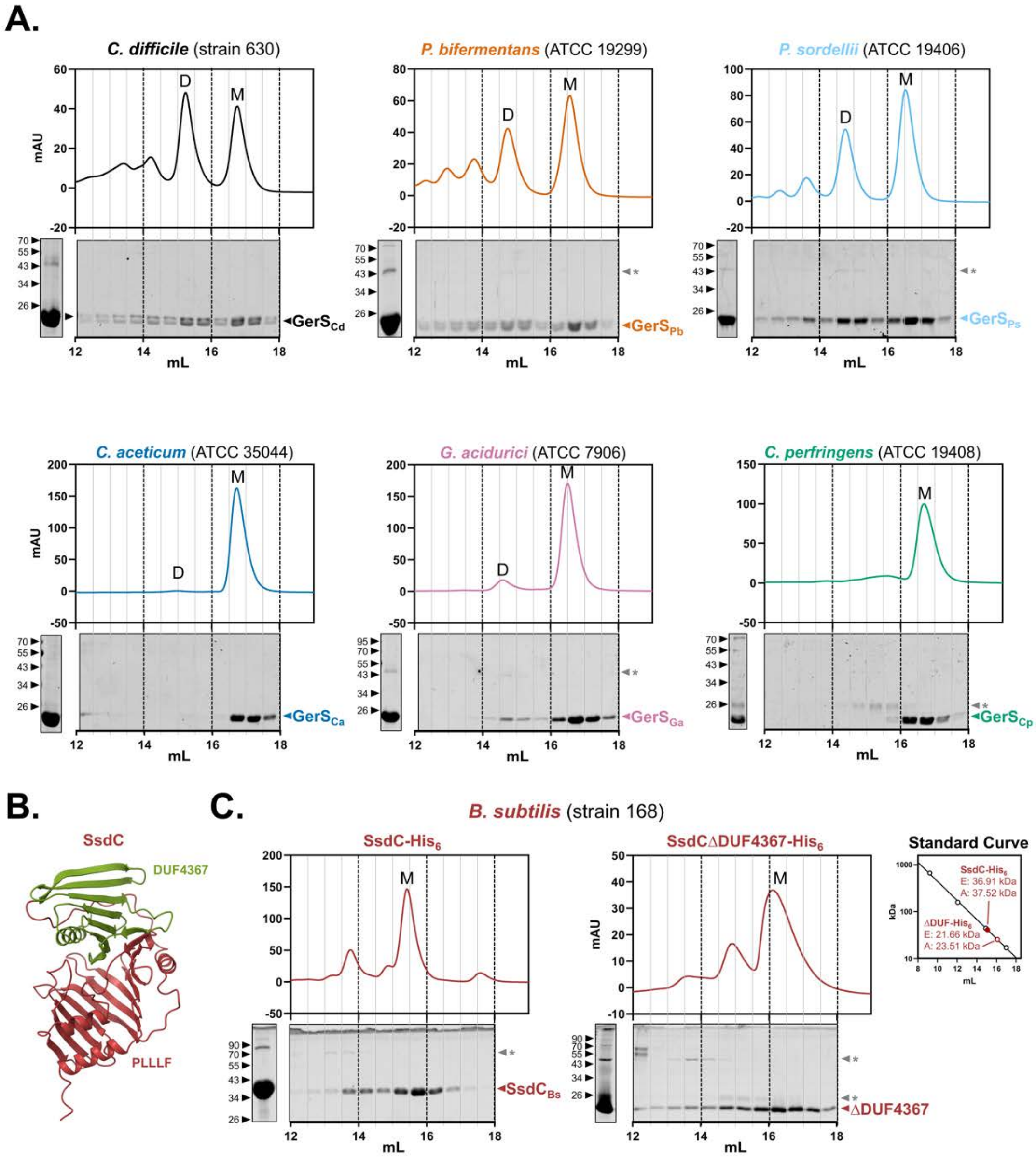


**Supplemental Figure 4. Fractions from SEC analyses of GerS orthologs.** A) Coomassie stain of fractions from SEC analyses of $His_6$-tagged GerS orthologs tag. Colored arrows point to GerS ortholog band. Gray asterisks mark likely *E. coli* chaperones that copurified with the protein. Coomassie stain of the input fractions analyzed by SEC are shown to the left of SEC fractions gels. MW is shown in kDa. B) AlphaFold3-predicted model [47] of SsdC with its DUF4367 domain. C) SEC analysis and Coomassie-stained fractions of $His_6$-tagged SsdC expressed with and without its DUF4367 domain. Apparent molecular (A) weights are plotted on the standard curve generated from SEC with Bio-Rad gel filtration standard and are compared with estimated molecular weights (E) calculated from the amino acid sequence. Layout is the same as described in panel A.

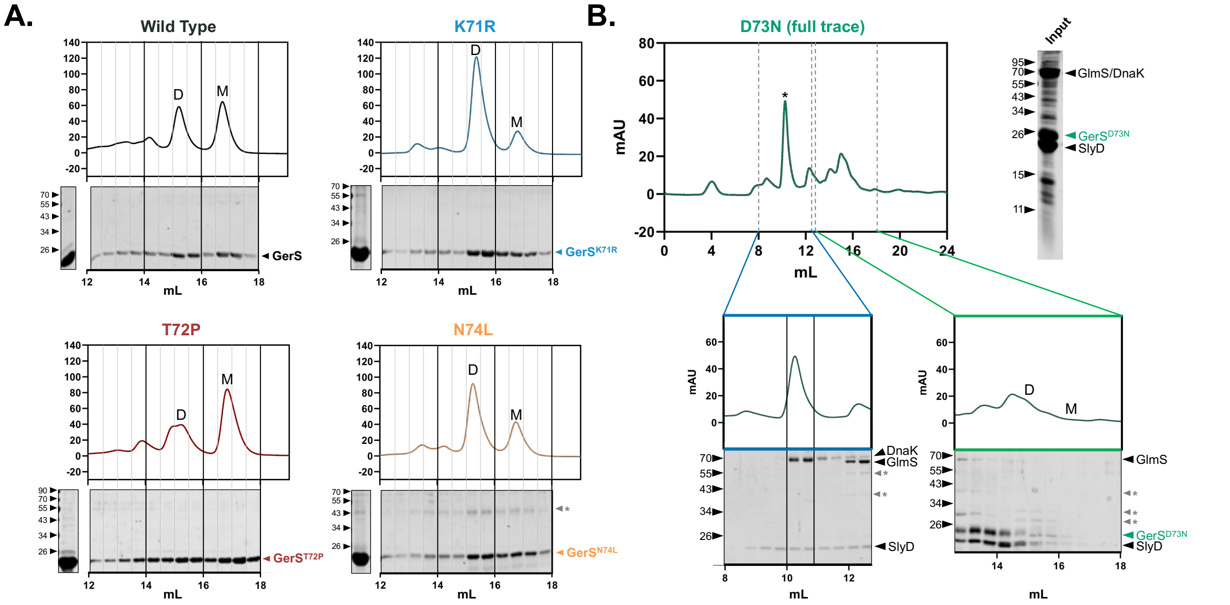


**Supplemental Figure 5. Fractions from SEC analyses of His$_6$-tagged GerS variants carrying LolA-guided hinge substitutions.** A) Coomassie stain of the SEC fractions. Colored arrows point to purified GerS orthologs. Gray asterisks mark likely *E. coli* chaperones that copurified with the GerS orthologs. Coomassie stains of the input fractions analyzed by SEC are shown to the left of SEC fraction gels. MW is shown in kDa. B) Full and partial SEC traces of GerS$^{D73N}$-His$_6$, with the Coomassie stain of the SEC input and fractions shown below. The black arrows point to *E. coli* chaperone proteins identified using Edman sequencing.

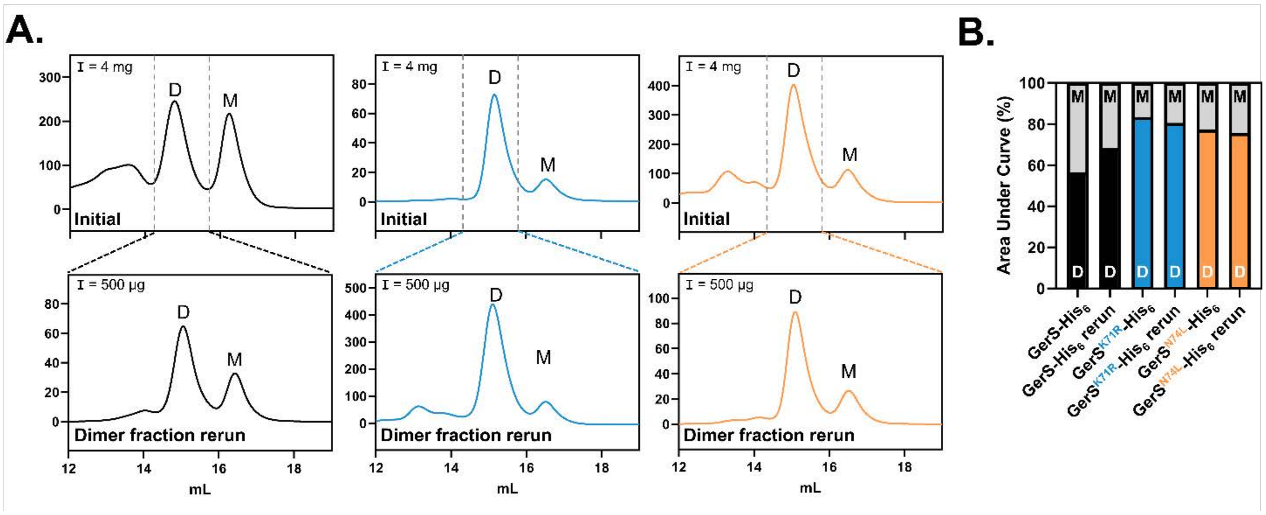


**Supplemental Figure 6. SEC analysis of affinity-purified GerS$^{K71R}$-His$_6$ and GerS$^{N74L}$-His$_6$.** A) Traces of the initial SEC analysis (top) and the rerun of the purified dimer fraction (bracketed by the vertical dashed lines) for GerS$^{WT}$-His$_6$, GerS$^{K71R}$-His$_6$, and GerS$^{N74L}$-His$_6$. I = input. The total amount of protein loaded is indicated in the top left of the traces. Solid lines represent the GerS variant, and dotted lines represent the fractions used in the rerun. B) Ratios of areas under each peak. The black column is the area under the dimer peak, and the gray column is the area under the monomer peak. Ratios are represented as a percentage of the total area of the monomer and dimer peaks combined.

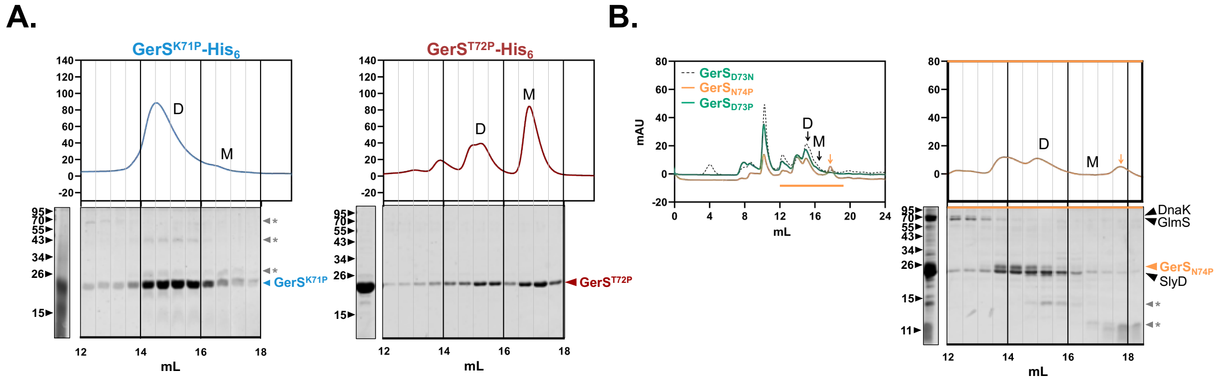


**Supplemental Figure 7. SEC traces and fractions of unstable GerS proline variants.** A) SEC traces of GerS$^{K71P}$-His$_6$ (left) and GerS$^{T72P}$-His$_6$ (right) with Coomassie stains of the fractions shown below. Black arrows correspond to non-GerS bands that co-purified with the His$_6$-tagged GerS variants. Traces are labeled with “D” for dimer or “M” for monomer based on the elution volumes for each peak. Gray asterisks mark likely *E. coli* chaperones that copurified with the protein. B) Full SEC traces of GerS$^{D73P}$-His$_6$ (green solid line) and GerS$^{N74P}$-His$_6$ (gold solid line) were aligned with the full SEC trace of GerS$^{D73N}$-His$_6$ (green dotted line). Gold arrow marks a peak that differs from GerS$^{D73P}$-His$_6$ and GerS$^{N74P}$-His$_6$ traces. Inset of GerS$^{N74P}$-His$_6$ with unique peak and Coomassie-stained fractions (right).

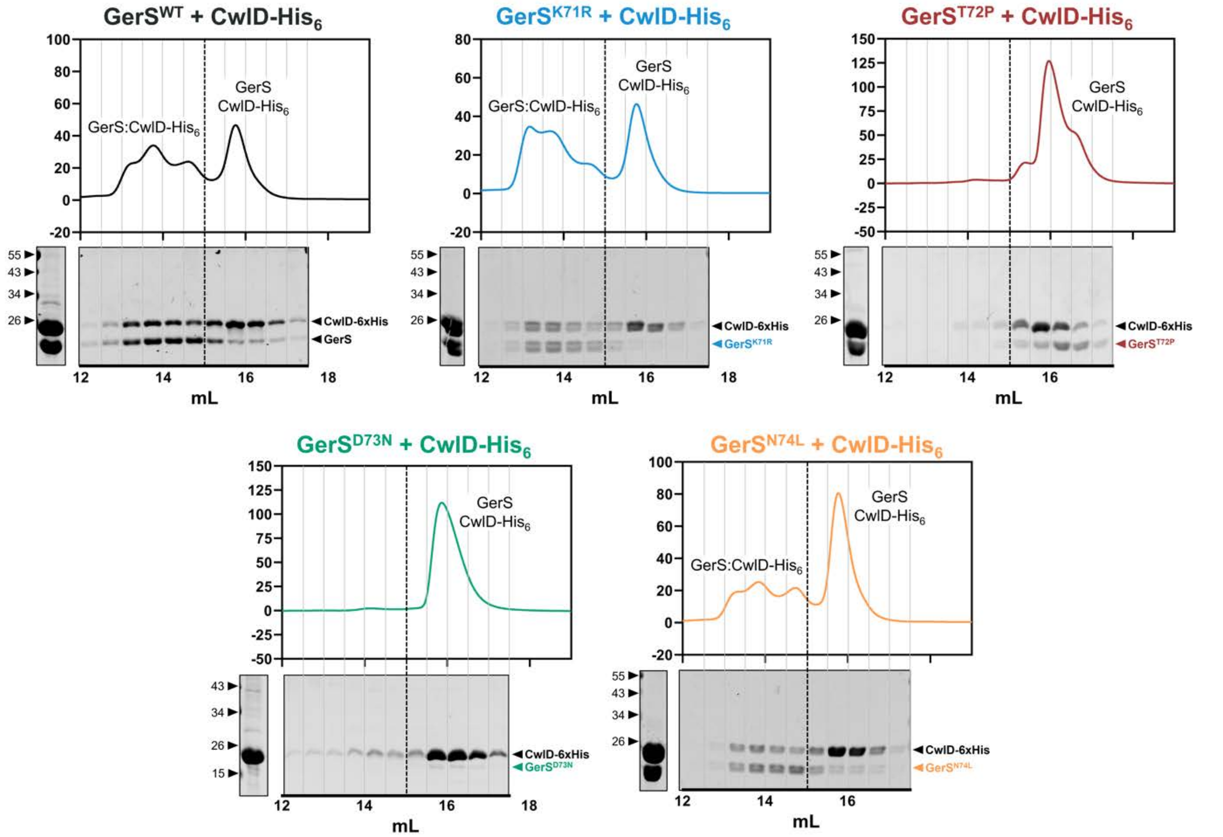


**Supplemental Figure 8. SEC analyses of untagged GerS variants carrying LolA-guided hinge substitutions following co-purification with CwlD-His$_6$.** SEC traces and Coomassie-stained fractions of untagged GerS hinge variants co-purified with His$_6$-tagged CwlD are shown. Coomassie-stained SEC inputs are shown to the left of each fraction gel. Peaks labeled with "GerS:CwlD-His6", correspond to the complex formed between GerS and CwlD-His$_6$. Peaks labeled with "GerS" and "CwlD-His$_6$", represents the proteins eluting outside of complex. MW is shown in kDa.

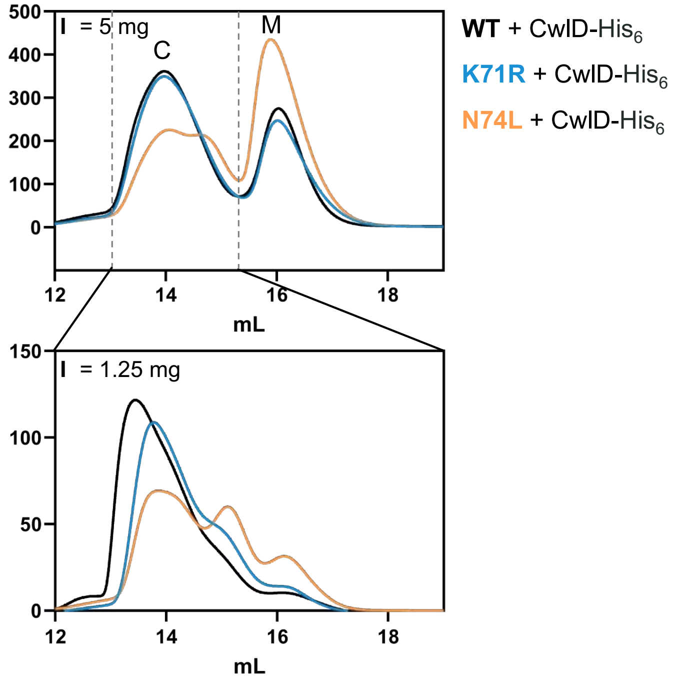


**Supplemental Figure 9. GerS homodimer-stabilizing mutations do not increase CwlD complex stability.** Aligned SEC traces of GerS$^{WT}$, GerS$^{K71R}$, and GerS$^{N74L}$ co-affinity purified with CwlD-His$_6$ (top) and SEC trace reruns of the GerS:CwlD complex fractions (bottom). Peaks are labeled "C" for GerS:CwlD-His$_6$ complex or "M" for monomers. Total protein loaded is shown in the top left of the traces.

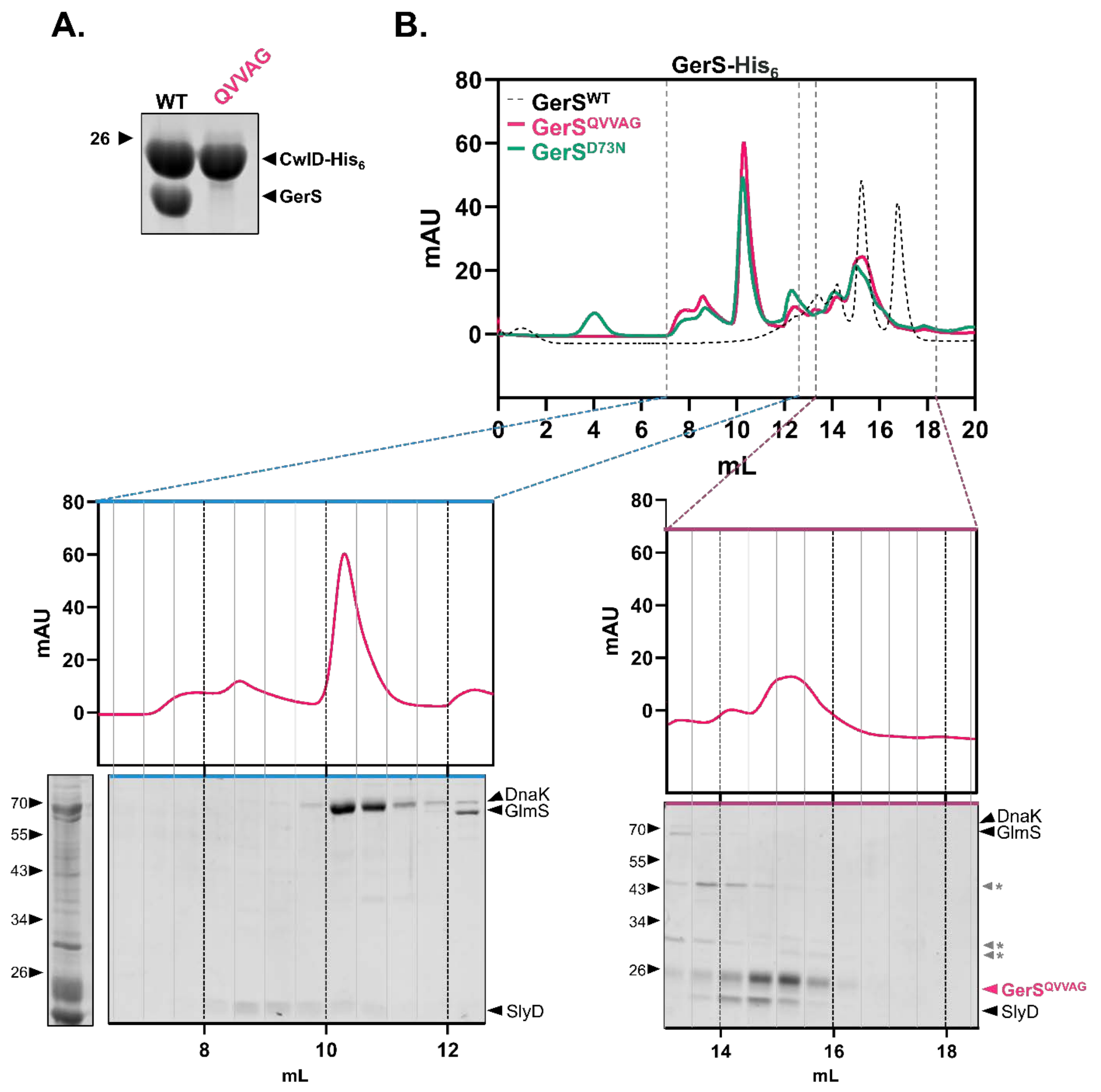


**Supplemental Figure 10. GerS$^{QVVAG}$-His$_6$ exhibits a similar SEC trace to the unstable GerS$^{D73N}$-His$_6$.** A) Coomassie stain of untagged GerS$^{QVVAG}$ variant co-purified with His$_6$-tagged CwlD. B) SEC trace of GerS$^{QVVAG}$-His$_6$ aligned with the SEC trace of GerS$^{D73N}$-His$_6$ and GerS$^{WT}$-His$_6$. Partial SEC traces of GerS$^{QVVAG}$-His$_6$ are shown along with fractions. Gray asterisks mark likely *E. coli* chaperones that copurified with the protein.

| % Identity | | | | | | | | | | | | | | | | | | |
|---|---|---|---|---|---|---|---|---|---|---|---|---|---|---|---|---|---|---|
| **Species** | **Locus Tag** | **1** | **2** | **3** | **4** | **5** | **6** | **7** | **8** | **9** | **10** | **11** | **12** | **13** | **14** | **15** | **16** | **17** |
| 1 - Acidobacterium_capsulatum_(ATCC_51196) | acp_3011 | | 6.7 | 9 | 10.6 | 10.6 | 7.2 | 13.2 | 7.9 | 5.9 | 11.7 | 9.3 | 15 | 8.9 | 7.5 | 6.4 | 13.1 | 7.8 |
| 2 - Bacillus_subtilis_(strain_168) | BSU04630 | 6.7 | | 7 | 9.4 | 9.8 | 21 | 8 | 8.9 | 5.7 | 10 | 9.9 | 9.6 | 11.9 | 12.8 | 7.7 | 8.7 | 7 |
| 3 - Bacteroides_uniformis_(ATCC_8492) | bacuni_04723 | 9 | 7 | | 16.1 | 13.8 | 7.5 | 19.6 | 10 | 7.9 | 11.5 | 11.4 | 12.6 | 10.6 | 9.4 | 8.9 | 12.1 | 7.4 |
| 4 - Bdellovibrio_bacteriovorus_(ATCC_15356) | bd3842 | 10.6 | 9.4 | 16.1 | | 13.5 | 10.8 | 14.3 | 9.7 | 16.2 | 16.8 | 11.5 | 15.1 | 9.4 | 13 | 7.9 | 13.1 | 6.3 |
| 5 - Borreliella_burgdorferi_(ATCC_35210) | bb_0346 | 10.6 | 9.8 | 13.8 | 13.5 | | 6.7 | 14.1 | 7.1 | 8.2 | 11.8 | 25.5 | 15.7 | 9.8 | 10 | 5.8 | 13.9 | 7.8 |
| 6 - Clostridioides_difficile_630 | CD630_34640 | 7.2 | 21 | 7.5 | 10.8 | 6.7 | | 10.4 | 8.6 | 4.2 | 7.9 | 7.1 | 10.1 | 12.9 | 7.9 | 5.2 | 7.4 | 5.9 |
| 7 - Flavobacterium_suncheonense | q764_01725 | 13.2 | 8 | 19.6 | 14.3 | 14.1 | 10.4 | | 6.1 | 10.3 | 13.4 | 12.2 | 16.2 | 8.3 | 10.8 | 6.8 | 14 | 8.2 |
| 8 - Fusobacterium_nucleatum_(ATCC_25586) | fn1493 | 7.9 | 8.9 | 10 | 9.7 | 7.1 | 8.6 | 6.1 | | 5.8 | 11.3 | 7.8 | 8.2 | 6.1 | 6.4 | 5.5 | 11.2 | 6.4 |
| 9 - Gloeobacter_kilaueensis_(ATCC_BAA-2537) | GKIL_3092 | 5.9 | 5.7 | 7.9 | 16.2 | 8.2 | 4.2 | 10.3 | 5.8 | | 9.1 | 8.7 | 10.8 | 4.2 | 8.4 | 7 | 8.5 | 3.4 |
| 10 - Helicobacter_mustelae_(ATCC_43772) | HMU08660 | 11.7 | 10 | 11.5 | 16.8 | 11.8 | 7.9 | 13.4 | 11.3 | 9.1 | | 12.7 | 13.5 | 8.8 | 14.3 | 8 | 15.7 | 7.3 |
| 11 - Leptospira_yanagawae_(ATCC_700523) | LEP1GSC202_2292 | 9.3 | 9.9 | 11.4 | 11.5 | 25.5 | 7.1 | 12.2 | 7.8 | 8.7 | 12.7 | | 13.5 | 9.9 | 10.1 | 10.5 | 19.7 | 9.5 |
| 12 - Myxococcus_fulvus_(ATCC_BAA-855) | LILAB_01365 | 15 | 9.6 | 12.6 | 15.1 | 15.7 | 10.1 | 16.2 | 8.2 | 10.8 | 13.5 | 13.5 | | 7.9 | 12.3 | 9.6 | 18.8 | 11.4 |
| 13 - Nitrosomonas_europaea_(ATCC_19718) | ne2245 | 8.9 | 11.9 | 10.6 | 9.4 | 9.8 | 12.9 | 8.3 | 6.1 | 4.2 | 8.8 | 9.9 | 7.9 | | 6.6 | 6.5 | 8.5 | 7.3 |
| 14 - Rickettsia_bellii | rbe_0046 | 7.5 | 12.8 | 9.4 | 13 | 10 | 7.9 | 10.8 | 6.4 | 8.4 | 14.3 | 10.1 | 12.3 | 6.6 | | 7.8 | 10.8 | 5.4 |
| 15 - Streptomyces_rapamycinicus_(ATCC_29253) | d3c57_120570 | 6.4 | 7.7 | 8.9 | 7.9 | 5.8 | 5.2 | 6.8 | 5.5 | 7 | 8 | 10.5 | 9.6 | 6.5 | 7.8 | | 9.9 | 5.5 |
| 16 - Escherchia coli (strain K12) | b0891 | 13.1 | 8.7 | 12.1 | 13.1 | 13.9 | 7.4 | 14 | 11.2 | 8.5 | 15.7 | 19.7 | 18.8 | 8.5 | 10.8 | 9.9 | | 11.2 |
| 17 - Mycobacterium tuberculosis | P9WK65 | 7.8 | 7 | 7.4 | 6.3 | 7.8 | 5.9 | 8.2 | 6.4 | 3.4 | 7.3 | 9.5 | 11.4 | 7.3 | 5.4 | 5.5 | 11.2 | |
| **% Similarity** | | | | | | | | | | | | | | | | | | |
| 1 - Acidobacterium_capsulatum_(ATCC_51196) | acp_3011 | | 36.6 | 41.9 | 46.8 | 41.3 | 38.7 | 48.6 | 31.5 | 33.8 | 38.3 | 38.7 | 48.2 | 33.9 | 41.2 | 28.5 | 38.8 | 29.9 |
| 2 - Bacillus_subtilis_(strain_168) | BSU04630 | 36.6 | | 34.3 | 41.8 | 35.6 | 55.2 | 41.8 | 32.6 | 29.9 | 35.2 | 37.1 | 37.7 | 40.7 | 39.3 | 26.1 | 36.5 | 27 |
| 3 - Bacteroides_uniformis_(ATCC_8492) | bacuni_04723 | 41.9 | 34.3 | | 43.9 | 40.1 | 33.5 | 63.3 | 31.4 | 32.8 | 37.3 | 45 | 45 | 37.5 | 42.9 | 27.9 | 37.9 | 30.5 |
| 4 - Bdellovibrio_bacteriovorus_(ATCC_15356) | bd3842 | 46.8 | 41.8 | 43.9 | | 41.9 | 39.2 | 55.2 | 35 | 43.1 | 41.6 | 40.4 | 48.3 | 36 | 48.1 | 32.6 | 46.8 | 27.7 |
| 5 - Borreliella_burgdorferi_(ATCC_35210) | bb_0346 | 41.3 | 35.6 | 40.1 | 41.9 | | 35.9 | 47.3 | 29.8 | 30.6 | 34.1 | 60 | 43 | 33.1 | 40 | 28.3 | 41.4 | 31.1 |
| 6 - Clostridioides_difficile_630 | CD630_34640 | 38.7 | 55.2 | 33.5 | 39.2 | 35.9 | | 45.8 | 30.6 | 26.8 | 31.5 | 35.1 | 34.9 | 42.2 | 36.1 | 25.9 | 34.9 | 28.3 |
| 7 - Flavobacterium_suncheonense | q764_01725 | 48.6 | 41.8 | 63.3 | 55.2 | 47.3 | 45.8 | | 32.9 | 36.9 | 41.7 | 49.8 | 49.8 | 39.4 | 50.2 | 32.7 | 43.7 | 32.5 |
| 8 - Fusobacterium_nucleatum_(ATCC_25586) | fn1493 | 31.5 | 32.6 | 31.4 | 35 | 29.8 | 30.6 | 32.9 | | 26 | 35.6 | 31.5 | 30.1 | 30.2 | 30.9 | 27.6 | 34.9 | 21.4 |
| 9 - Gloeobacter_kilaueensis_(ATCC_BAA-2537) | GKIL_3092 | 33.8 | 29.9 | 32.8 | 43.1 | 30.6 | 26.8 | 36.9 | 26 | | 28.1 | 30.7 | 35.7 | 23.3 | 34.7 | 27.2 | 29.3 | 19.7 |
| 10 - Helicobacter_mustelae_(ATCC_43772) | HMU08660 | 38.3 | 35.2 | 37.3 | 41.6 | 34.1 | 31.5 | 41.7 | 35.6 | 28.1 | | 37.3 | 33.6 | 28.5 | 37.8 | 27.7 | 41.3 | 22.3 |
| 11 - Leptospira_yanagawae_(ATCC_700523) | LEP1GSC202_2292 | 38.7 | 37.1 | 45 | 40.4 | 60 | 35.1 | 49.8 | 31.5 | 30.7 | 37.3 | | 40.2 | 29.7 | 42 | 30.5 | 43.1 | 27 |
| 12 - Myxococcus_fulvus_(ATCC_BAA-855) | LILAB_01365 | 48.2 | 37.7 | 45 | 48.3 | 43 | 34.9 | 49.8 | 30.1 | 35.7 | 33.6 | 40.2 | | 33.4 | 41.5 | 31.4 | 57.6 | 39.1 |
| 13 - Nitrosomonas_europaea_(ATCC_19718) | ne2245 | 33.9 | 40.7 | 37.5 | 36 | 33.1 | 42.2 | 39.4 | 30.2 | 23.3 | 28.5 | 29.7 | 33.4 | | 33.9 | 24 | 30.3 | 23.3 |
| 14 - Rickettsia_bellii | rbe_0046 | 41.2 | 39.3 | 42.9 | 48.1 | 40 | 36.1 | 50.2 | 30.9 | 34.7 | 37.8 | 42 | 41.5 | 33.9 | | 27.4 | 39.4 | 23.8 |
| 15 - Streptomyces_rapamycinicus_(ATCC_29253) | d3c57_120570 | 28.5 | 26.1 | 27.9 | 32.6 | 28.3 | 25.9 | 32.7 | 27.6 | 27.2 | 27.7 | 30.5 | 31.4 | 24 | 27.4 | | 33.9 | 21.6 |
| 16 - Escherchia coli (strain K12) | b0891 | 38.8 | 36.5 | 37.9 | 46.8 | 41.4 | 34.9 | 43.7 | 34.9 | 29.3 | 41.3 | 43.1 | 57.6 | 30.3 | 39.4 | 33.9 | | 35.2 |
| 17 - Mycobacterium tuberculosis | P9WK65 | 29.9 | 27 | 30.5 | 27.7 | 31.1 | 28.3 | 32.5 | 21.4 | 19.7 | 22.3 | 27 | 39.1 | 23.3 | 23.8 | 21.6 | 35.2 | |

**Table S1. Percent identity and percent similarity calculations from aligned PLLLF proteins.**

| KBase Reference | Genome Object Name |
|---|---|
| 201601/10/1 | *Caulobacter_vibrioides* |
| 201601/169/1 | *Acidithrix_ferrooxidans* |
| 201601/90/1 | *Blautia_faecis* |
| 201601/238/1 | *Limihaloglobus_sulfuriphilus* |
| 201601/272/1 | *Solitalea_canadensis_DSM_3403* |
| 201601/2/1 | *Bacillus_subtilis_subsp._subtilis_str._168* |
| 201601/112/1 | *Aminomonas_paucivorans_DSM_12260* |
| 201601/160/1 | *Denitrobacterium_detoxificans* |
| 201601/24/1 | *Akkermansia_glycaniphila* |
| 201601/241/1 | *Criblamydia_sequanensis_CRIB-18* |
| 201601/242/1 | *Estrella_lausannensis* |
| 201601/243/1 | *Parachlamydia_acanthamoebae* |
| 201601/5/1 | *Mycoplasma_agalactiae* |
| 201601/40/1 | *Coprothermobacter_platensis_DSM_11748* |
| 201601/244/1 | *Waddlia_chondrophila_WSU_86-1044* |
| 201601/247/1 | *Victivallis_vadensis* |
| 201601/25/1 | *Chlamydia_abortus* |
| 201601/200/1 | *Agrobacterium_fabrum_str._C58* |
| 201601/250/1 | *Pedosphaera_parvula_Ellin514* |
| 201601/252/1 | *Brevifollis_gellanilyticus* |
| 201601/273/1 | *Sphingobacterium_mizutaii_NBRC_14946___DSM_11724* |
| 201601/102/1 | *Geovibrio_sp._L21-Ace-BES* |
| 201601/253/1 | *Limisphaera_ngatamarikiensis* |
| 201601/276/1 | *Roseiflexus_castenholzii_DSM_13941* |
| 201601/91/1 | *Butyricicoccus_pullicaecorum* |
| 201601/277/1 | *Oscillochloris_trichoides_DG-6* |
| 201601/119/1 | *Kosmotoga_arenicorallina_S304* |
| 201601/113/1 | *Cloacibacillus_evryensis_DSM_19522* |
| 201601/278/1 | *Herpetosiphon_llansteffanensis* |
| 201601/158/1 | *Atopobium_rimae_ATCC_49626* |
| 201601/279/1 | *Chloroflexus_islandicus* |
| 201601/28/1 | *Acanthopleuribacter_pedis* |
| 201601/280/1 | *Kallotenue_sp._CFH_73958* |
| 201601/115/1 | *Fretibacterium_fastidiosum* |
| 201601/161/1 | *Cryptobacterium_curtum_DSM_15641* |
| 201601/107/1 | *Desulfurispirillum_indicum_S5* |
| 201601/50/1 | *Tissierella_creatinophila_DSM_6911* |
| 201601/175/1 | *Streptomyces_albofaciens_JCM_4342* |
| 201601/254/1 | *Rubritalea_marina_DSM_17716* |
| 201601/32/1 | *Leptospirillum_ferriphilum* |
| 201601/33/1 | *Fervidobacterium_changbaicum* |
| 201601/201/1 | *Rickettsia_aeschlimannii* |
| 201601/34/1 | *Geotoga_petraea* |
| 201601/36/1 | *Acetomicrobium_hydrogeniformans_ATCC_BAA-1850* |
| 201601/38/1 | *Balnearium_lithotrophicum* |
| 201601/51/1 | *Adlercreutzia_equolifaciens_DSM_19450* |
| 201601/94/1 | *Aquifex_aeolicus_VF5* |
| 201601/53/1 | *Acidimicrobium_ferrooxidans_DSM_10331* |

| | |
|---|---|
| 201601/26/1 | *Lentisphaera_araneosa_HTCC2155* |
| 201601/118/1 | *Athalassotoga_saccharophila* |
| 201601/56/1 | *Acholeplasma_axanthum* |
| 201601/57/1 | *Entomoplasma_freundtii* |
| 201601/58/1 | *Mesoplasma_coleopterae* |
| 201601/202/1 | *Rhodobacter_blasticus_DSM_2131* |
| 201601/170/1 | *Desertimonas_flava* |
| 201601/59/1 | *Spiroplasma_atrichopogonis* |
| 201601/6/1 | *Anaerolinea_thermolimosa* |
| 201601/95/1 | *Persephonella_atlantica* |
| 201601/62/1 | *Abyssicoccus_albus* |
| 201601/63/1 | *Aerococcus_urinae* |
| 201601/12/1 | *Myxococcus_fulvus* |
| 201601/64/1 | *Anoxybacillus_flavithermus* |
| 201601/39/1 | *Deferribacter_autotrophicus* |
| 201601/65/1 | *Lacticaseibacillus_paracasei* |
| 201601/260/1 | *Chloroherpeton_thalassium_ATCC_35110* |
| 201601/203/1 | *Azospirillum_agricola* |
| 201601/68/1 | *Acetonema_longum_DSM_6540* |
| 201601/269/1 | *Arcticibacter_eurypsychrophilus* |
| 201601/96/1 | *Thermocrinis_minervae* |
| 201601/7/1 | *Deinococcus_actinosclerus* |
| 201601/70/1 | *Pectinatus_frisingensis* |
| 201601/71/1 | *Selenomonas_artemidis_DSM_19719* |
| 201601/8/1 | *Armatimonas_rosea* |
| 201601/80/1 | *Anaerococcus_degeneri* |
| 201601/4/1 | *Synechococcus_elongatus_PCC_11802* |
| 201601/81/1 | *Finegoldia_magna* |
| 201601/120/1 | *Marinitoga_lauensis* |
| 201601/123/1 | *Dictyoglomus_turgidum_DSM_6724* |
| 201601/264/1 | *Alkaliflexus_imshenetskii_DSM_15055* |
| 201601/172/1 | *Ilumatobacter_sp._SYSU_D60003* |
| 201601/206/1 | *Chondromyces_apiculatus_DSM_436* |
| 201601/97/1 | *Phorcysia_thermohydrogeniphila* |
| 201601/128/1 | *Coprothermobacter_proteolyticus_DSM_5265* |
| 201601/13/1 | *Helicobacter_pylori* |
| 201601/41/1 | *Caldisericum_exile_AZM16c01* |
| 201601/270/1 | *Mucilaginibacter_galii* |
| 201601/100/1 | *Calditerrivibrio_nitroreducens_DSM_19672* |
| 201601/131/1 | *Capsulimonas_corticalis* |
| 201601/135/1 | *Aliterella_atlantica_CENA595* |
| 201601/136/1 | *Anabaena_aphanizomenioides_LEGE_00250* |
| 201601/137/1 | *Microcystis_aeruginosa_KW* |
| 201601/82/1 | *Lagierella_massiliensis* |
| 201601/138/1 | *Spirulina_major_PCC_6313* |
| 201601/14/1 | *Bdellovibrio_bacteriovorus* |
| 201601/174/1 | *Frankia_asymbiotica* |
| 201601/265/1 | *Dysgonomonas_capnocytophagoides_DSM_22835* |
| 201601/142/1 | *Pelolinea_submarina* |

| | |
|---|---|
| 201601/48/1 | *Clostridioides_difficile_630* |
| 201601/259/1 | *Prosthecochloris_vibrioformis* |
| 201601/143/1 | *Ornatilinea_apprima* |
| 201601/144/1 | *Levilinea_saccharolytica* |
| 201601/145/1 | *Bellilinea_caldifistulae_1* |
| 201601/263/1 | *Alistipes_communis* |
| 201601/101/1 | *Flexistipes_sinusarabici_DSM_4947* |
| 201601/146/1 | *Calidithermus_chliarophilus_DSM_9957* |
| 201601/149/1 | *Marinithermus_hydrothermalis_DSM_14884* |
| 201601/15/1 | *Mariprofundus_aestuarium* |
| 201601/150/1 | *Oceanithermus_profundus_DSM_14977* |
| 201601/175/1 | *Streptomyces_albofaciens_JCM_4342* |
| 201601/208/1 | *Vulgatibacter_incomptus* |
| 201601/151/1 | *Thermus_islandicus_DSM_21543* |
| 201601/42/1 | *Chrysiogenes_arsenatis_DSM_11915* |
| 201601/176/1 | *Corynebacterium_accolens_ATCC_49725* |
| 201601/83/1 | *Peptoniphilus_grossensis* |
| 201601/177/1 | *Arsenicicoccus_bolidensis_DSM_15745* |
| 201601/266/1 | *Salinivirga_cyanobacteriivorans* |
| 201601/103/1 | *Seleniivibrio_woodruffii* |
| 201601/18/1 | *Bacteroides_pyogenes* |
| 201601/180/1 | *Aliiarcobacter_faecis* |
| 201601/181/1 | *Arcobacter_butzleri* |
| 201601/27/1 | *Borrelia_burgdorferi_IPT24* |
| 201601/209/1 | *Archangium_violaceum* |
| 201601/182/1 | *Campylobacter_armoricus* |
| 201601/183/1 | *Nitratiruptor_tergarcus_DSM_16512* |
| 201601/171/1 | *Ferrimicrobium_acidiphilum_DSM_19497* |
| 201601/43/1 | *Caldithrix_abyssi_DSM_13497* |
| 201601/186/1 | *Alkalispirochaeta_alkalica_DSM_8900* |
| 201601/187/1 | *Brachyspira_hyodysenteriae* |
| 201601/188/1 | *Leptonema_illini_DSM_21528* |
| 201601/154/1 | *Gemmatimonas_aurantiaca_T-27* |
| 201601/189/1 | *Spirochaeta_cellobiosiphila_DSM_17781* |
| 201601/19/1 | *Chlorobaculum_limnaeum* |
| 201601/106/1 | *Desulfurispira_natronophila* |
| 201601/21/1 | *Gemmatirosa_kalamazoonesis* |
| 201601/195/1 | *Geothrix_fermentans_DSM_14018* |
| 201601/88/1 | *Clostridium_perfringens* |
| 201601/114/1 | *Jonquetella_anthropi_DSM_22815* |
| 201601/196/1 | *Holophaga_foetida_DSM_6591* |
| 201601/324/1 | *Fusobacterium_varium_ATCC_27725* |
| 201601/319/1 | *Fusobacterium_gonidiaformans_ATCC_25563* |
| 201601/322/1 | *Psychrilyobacter_atlanticus_DSM_19335* |
| 201601/321/1 | *Streptobacillus_moniliformis_DSM_12112* |
| 201601/320/1 | *Leptotrichia_goodfellowii_DSM_19756* |
| 201601/159/1 | *Collinsella_aerofaciens* |
| 201601/212/1 | *Ghiorsea_bivora* |
| 201601/216/1 | *Thermithiobacillus_tepidarius_DSM_3134* |

| | |
|---|---|
| 201601/89/1 | *Acetivibrio_ethanolgignens* |
| 201601/44/1 | *Dictyoglomus_thermophilum* |
| 201601/219/1 | *Hydrogenophilus_thermoluteolus* |
| 201601/222/1 | *Vibrio_alginolyticus_NBRC_15630___ATCC_17749* |
| 201601/223/1 | *Acinetobacter_baumannii* |
| 201601/224/1 | *Shewanella_baltica* |
| 201601/17/1 | *Tepidiphilus_margaritifer_DSM_15129* |
| 201601/11/1 | *Neisseria_gonorrhoeae* |
| 201601/271/1 | *Parapedobacter_koreensis* |
| 201601/225/1 | *Oceanococcus_atlanticus* |
| 201601/229/1 | *Acidovorax_carolinensis* |
| 201601/9/1 | *Escherichia_coli* |
| 201601/16/1 | *Acidithiobacillus_caldus_ATCC_51756* |
| 201601/197/1 | *Thermotomaculum_hydrothermale* |
| 201601/23/1 | *Algisphaera_agarilytica* |
| 201601/230/1 | *Bordetella_avium* |
| 201601/231/1 | *Oxalobacter_formigenes* |
| 201601/232/1 | *Methylophilus_methylotrophus_DSM_46235___ATCC_53528* |
| 201601/235/1 | *Sedimentisphaera_salicampi* |
| 201601/49/1 | *Veillonella_atypica* |
| 201601/236/1 | *Phycisphaerae_bacterium_RAS1* |
| 201601/237/1 | *Anaerohalosphaera_lusitana* |

**Table S2. List of KBase genomes used to generate phylogenetic tree.**
